# An endogenous peptide PEP2 modulates Iron-deficiency signalling and root growth in Arabidopsis

**DOI:** 10.1101/2024.09.10.612258

**Authors:** Deep Shikha, Sanskar Mishra, Santosh B. Satbhai

## Abstract

Iron (Fe) is an essential element for most of the living organisms including plants and humans, where plants serve as the primary source of our dietary iron intake. The availability of Fe determines plant fitness and yield. Thus, understanding of iron uptake, its acquisition and utilization is of critical importance to reap nutritional benefits from plant breeding. Despite significant progress in uncovering how iron homeostasis is regulated by transcription factors and phytohormones, molecular pathways that mediate Fe deficiency through the action of signalling peptides remain elusive. In this work, we reported the role of PROPEP2 (Plant Elicitor Peptide 2) and PEPR (Perception of Arabidopsis danger signal peptide receptor) in regulating plant growth and development under Fe-deprived conditions. We revealed that a Damage Associated Molecular Pattern (DAMP) such as PROPEP2 is significantly induced under Fe deficiency. We also show that PEP2 modulates the expression of Iron Regulated Transporter 1 (IRT1) and Ferric-Reduction Oxidase (FRO2) under Fe deficiency. Furthermore, we showed that PEPR2 perceives PEP2 to positively regulate reactive oxygen species (ROS) content and negatively regulate the primary root growth, iron content and rhizosphere acidification. Our findings reveal the complex interplay between Fe and DAMP signalling pathways in plants.

## INTRODUCTION

Iron (Fe) is an essential micronutrient for virtually all forms of life, including plants. It acts as a cofactor of various enzymes like peroxidase (POD), ascorbate peroxidase (APX), and catalase (CAT) (Marschner & Römheld, 1994). Iron is required for diverse cellular and physiological processes that includes, cellular respiration, oxygen transport, chloroplast development, chlorophyll biosynthesis, DNA repair and photosynthesis in plants (Kobayashi & Nishizawa, 2012a; W. Li & Lan, 2017). It is the fourth most abundant mineral in earth’s crust and sixth most abundant element in the universe, yet most of it is inaccessible to plant utilization, due to its differential chemical properties in the oxygenated environment (Morgan & Anders, 1980). Iron bioavailability is also determined by various soil-dependent factors including soil microbiota, level of iron storage proteins, extent of iron-solubilization by redox reactions and level of synthetic and natural iron chelators. Approximately 30% of arable land is calcareous, which makes insoluble crystals of ferric (Fe^3+^) oxyhydrates unavailable for plants (Curie Catherine, 2003). Limited Fe availability induces diverse iron acquisition mechanisms in plants, which further induce changes at morphological, physiological, cellular, molecular, metabolic and gene expression levels to combat iron deficiency (Abadía et al., 2011; Jeong & Guerinot, 2009).

For efficient iron uptake from soil, plants adopt two distinct strategies: strategy □ (reduction-based strategy) and strategy □ (chelation-based strategy) (Brumbarova et al., 2015; Kobayashi & Nishizawa, 2012a). Non-graminaceous plants, including model plant *Arabidopsis thaliana* (Arabidopsis) use strategy □ to combat iron deficiency stress (Brumbarova et al., 2015; Kobayashi & Nishizawa, 2012a). On the other hand, graminaceous plants use strategy □, which involves secretion of mugineic acid (MA) family Phytosiderophores (MAs) as Fe^3+^-solubilizing molecules (Kobayashi & Nishizawa, 2012a; Marschner & Römheld, 1994). Thus far, nine types of MAs have been identified whose biosynthetic pathways are almost conserved (Kobayashi & Nishizawa, 2012a; Marschner & Römheld, 1994). These MA-Phytosiderophores (PS) are released by roots and form stable Fe-PS chelates, which are then taken up into root cells by Yellow-Strip 1 (YS1) and Yellow-Strip 1-Like (YS1L) plasma membrane (PM)-localized transporters (Römheld & Marschner, 1986). On the contrary, strategy □ response involves two steps. In the first step, rhizosphere is acidified via PM-localized -H+-ATPases which are encoded by Autoinhibited-H+-ATPases (AHA) (Santi & Schmidt, 2009). This acidification enhances Fe^3+^ solubility in the rhizosphere soil. Additional chelation and mobilization of Fe^3+^ is done by coumarin family phenolics which are released into the rhizosphere by the ABC transporter, PDR9/ABCG37 (Stringlis et al., 2019). Subsequently, chelated Fe^3+^ (insoluble form) is reduced to Fe^2+^ (soluble form) via Ferric-Reduction Oxidase (FRO2), a PM-localized ferric chelate reductase enzyme (Robinson et al., 1999). Soluble Fe^2+^ is then taken up into root cells by Iron Regulated Transporter 1 (IRT1) (Eide et al., 1996).

Under iron deficiency, many essential Fe uptake genes are induced to facilitate Fe uptake from rhizosphere. In Arabidopsis, FER-like iron deficiency-induced transcription factor (FIT), is induced by Fe deficiency and forms a heterodimer with subgroup Ib basic helix–loop–helix (bHLH) transcription factors (TFs) including *bHLH38*, *bHLH39*, *bHLH100* and *bHLH101* to further activate the transcription of *FRO2* and *IRT1* (Liang et al., 2017; Wang et al., 2013; Yuan et al., 2008). The expression of subgroup Ib bHLH TFs is induced by Fe deficiency, and this induction is further enhanced by subgroup IVc bHLH TFs including *bHLH115*, *bHLH104*, *ILR3/IAA-LEUCINE RESISTANT3 (bHLH105)*, and *bHLH34* (X. Li et al., 2016; Liang et al., 2017). ILR3, a member of subgroup IVc bHLH TF, has a dual role as transcriptional activator and repressor. It interacts with POPEYE (PYE) to down-regulate Fe distribution genes *(NAS4/NICOTIANAMINE SYNTHASE4, ZINC-INDUCED FACILITATOR1 (ZIF1)* and *FRO3*), storage genes (*FER1, FER3* and *FER4*) and assimilation genes (*NEET*) (Tissot et al., 2019). The activity of subgroup IVc genes is regulated post-transcriptionally to induce the expression of subgroup Ib genes only under Fe deficiency. BRUTUS (BTS), an E3 ubiquitin ligase, implicates this regulation by degrading subgroup IVc bHLH TFs under Fe-sufficient conditions (Selote et al., 2015). In addition to BTS, two closely related RING E3 ubiquitin ligases including BRUTUS LIKE1 (BTSL1) and BTSL2, directly target FIT for 26S proteasomal degradation to negatively regulate Fe homeostasis (Hindt et al., 2017). BTS also regulate Fe transport via FE UPTAKE-INDUCING PEPTIDE3 (FEP3)/IRON MAN1 (IMA1), a class of potential phloem-mobile small proteins that mediate iron transport systemically (Y. Li et al., 2021; Lichtblau et al., 2022).

Increasing evidence indicates that phytohormones including auxin indole-3-acetic acid (IAA), ethylene (ETH) and nitric oxide (NO), and ROS regulate Fe deficiency-induced adaptive responses in roots (Wolfgang Schmidt & Bartels, 1996; Sun et al., 2016; Wu et al., 2012; Zhai et al., 2016). Recently, a novel connection was uncovered between Fe homeostasis and plant immunity. Cao et al. have reported that FLG22/Flagellin, a MAMP/Microorganism Associated Molecular Pattern, degrades IMA1 to suppress root iron acquisition (Cao et al., 2024). However, how the perception of DAMP/ Damage Associated Molecular Patterns re-channelizes Fe signaling to trigger immune pathways downstream that determine the nature of plant responses remains unknown. Our study has attempted to get an insight into the molecular mechanism of crosstalk between Fe homeostasis and DAMP signalling.

DAMPs unlike microbe/pathogen associated molecular patterns (MAMPs/PAMPs), are endogenous molecules of the host (Boller & Felix, 2009). In Arabidopsis, several cytokine-like peptides, AtPeps, were initially identified as DAMPs (Boller & Felix, 2009). They belong to a small family of eight peptides (AtPep1-8) which originated from their respective precursors (AtPROPEP1-8) via metacaspase4 mediated cleavage at their respective C’ terminal portions (Handler, 2016). Handler et al., (2016) showed that in a non-damaged cell, inactive forms of metacaspases form complexes with PROPEPs, which reside in the cytosol, adjacent to the vacuolar tonoplast. Upon cell damage, Ca^2+^ influx activates metacaspases which further lead to modification of precursor PROPEPS to active PEPs upon cleavage. These PEPs are then exported extracellularly via an unconventional secretion system (Handler, 2016). Extracellularly, the PEPs are perceived by two homologous leucine –rich repeat (LRR) receptor-like kinases (RLK) i.e., PEPR1 and PEPR2 (Krol et al., 2010), which are encoded with an extracellular LRR domain and an intracellular protein kinase domain which triggers the downstream signalling upon their interaction with a co-receptor, Brassinosteroid Insensitive1-Associated Kinase1 (BAK1) (Krol et al., 2010). Interestingly, AtPep1 and AtPep2 are perceived by PEPR2 exclusively, in contrast, PEPR1 shows a broad-spectrum recognition for all AtPeps in a dose dependent manner (Yamaguchi et al., 2010). Upon receptor-ligand perception, a series of downstream events take place which include ion fluxes via plasma membrane localized ATPases, RbohD protein activation, ROS production, the release of volatile compounds as well as secondary metabolites, mitogen-activated protein kinase (MAPK)-phosphorylation via MAP Kinase cascades, phytohormones-responsive genes activation (including ethylene, jasmonic acid, and salicylic acid), transcriptional reprogramming of host cell and subsequent defense responses (Baxter et al., 2014; Yidong Liu & Zhang, 2004; Qia et al., 2010; Ranf et al., 2011; Rasmussen et al., 2012).

Several secreted peptides, including DAMPS are known to play a crucial role in regulating plant development similar to phytohormones although, molecular understanding in imparting plant stress responses, especially under nutrient stress conditions remains fragmentarily understood. Takashi et al., (2018) identified a peptide gene FEP1 (Fe-Uptake-Inducing Peptide1) that is induced under iron deficiency (Hirayama et al., 2018). CEP-1 (C-Terminally Encoded Peptide-1), a root to shoot signalling molecule, plays a crucial role in regulating lateral root formation and root nodulation under conditions of N-starvation (Aggarwal et al., 2020). Recently, a peptide signal relay between the small peptides OsPep3 and OsPSK have been studied in rice in response to wounding (Harshith et al., 2024). Likewise, AtPEP3 has been shown to regulate immune response and salinity stress tolerance in Arabidopsis (Nakaminami et al., 2018). Interestingly, Alanis et al. (2017) have reported the perception of CLE14/ CLAVATA3/ENDOSPERM SURROUNDING REGION peptide via CLV2/CLAVATA2 and PEPR2 receptors under phosphorous deficiency stress which triggers Fe mobilization in roots (Gutiérrez-Alanís et al., 2017). It is also known that PEP2 modulates the expression of CPC/CAPRICE and GL2/GLABRA2 to influence root hair cell fate in Arabidopsis (Jing et al., 2024).

Studies on PEP/PEPR signalling in controlled conditions have contributed to our knowledge on the crosstalk in plant development and immunity. However, the mechanisms by which PEP peptides are involved in modulating stress responses due to Fe deficiency remain unknown. In this study, we elucidated the role of the PEP2-PEPR2 module in maintaining the balance between Fe homeostasis and growth in Arabidopsis. We identified that limitation in Fe condition transcriptionally activates PEP2, to regulate the expression of genes such as, *IRT1*, *FRO2*, and *BTS*, which are involved in maintaining Fe homeostasis. Our work demonstrates that in response to Fe starvation, PEP2 is perceived by PEPR2 to positively regulate ROS and negatively regulate primary root growth, iron content and rhizosphere acidification. These findings suggest a possible overlap between Fe deficiency response signalling and PEP-PEPR signalling pathways in Arabidopsis.

## RESULTS

### Fe Deficiency Induces *PEP2* Expression

It is well known that promoter activity of *AtPROPEP1*, *AtPROPEP2* and *AtPROPEP3* is strongly activated by elf18, flg22, AtPeps, ethylene, MeJA, and wounding (Bartels & Boller, 2015; Safaeizadeh & Boller, 2019). Since, PROPEPs are induced upon cell damage and converted to mature PEPs in response to biotic as well as abiotic stress conditions, we thought to investigate their role in response to Fe limitation (Baez et al., 2022; Eliza P. Loo et al., 2020; Hou et al., 2019; Q. Li et al., 2020; Nakaminami et al., 2018). To assist in the visualization of cell damage of root meristematic zone (MZ) under Fe deficiency, we stained the wild type (WT/*Col-0*) roots with propidium iodide (PI), which stains cell walls as well as dead cells which show PM disruption and is impermeable through the PM of living cells. We found that epidermal cells in the MZ of WT roots were profusely stained at 72 hours of Fe deficiency and PI diffusion intracellularly indicates loss of cell vitality (Figure 1a). We further examined the status of cell viability using Evans blue staining and found an intense blue stain in the meristematic as well as elongation zone of WT at 72 hours of Fe deficiency indicating the loss of viability in the stained cells (Figure 1b). This loss in cell vitality under Fe deficiency may induce the expression of PROPEPs. To test this idea, we measured the expression levels of PROPEPs (1-6) at 72 hours of Fe deficiency using Arabidopsis eGFP browser (Figure S1a). This expression data suggests that *PROPEP2*, *PROPEP3* and *PROPEP5* are transcriptionally upregulated at significant levels in root tissues under iron-deficient conditions, with *PROPEP2* showing the highest expression among others (Figure S1a). Hence, we selected *PROPEP2*, as a candidate PEP showing elevated expression in epidermis and stele of root under Fe starvation (Figure S1b). To confirm this expression profile from the eGFP browser, we generated *pPROPEP2:GUS*/*Col-0* expressing lines and looked at spatial expression patterns of *PROPEP2* at the tissue level. We found a higher promoter activity of *PROPEP2* in a tissue-specific manner in stele of lateral root, maturation zone, elongation zone and root tip under iron-deficient conditions (Figure 1c). These results confirm the transcriptional induction of *PROPEP2* under Fe deficient conditions.

**Figure 1.**
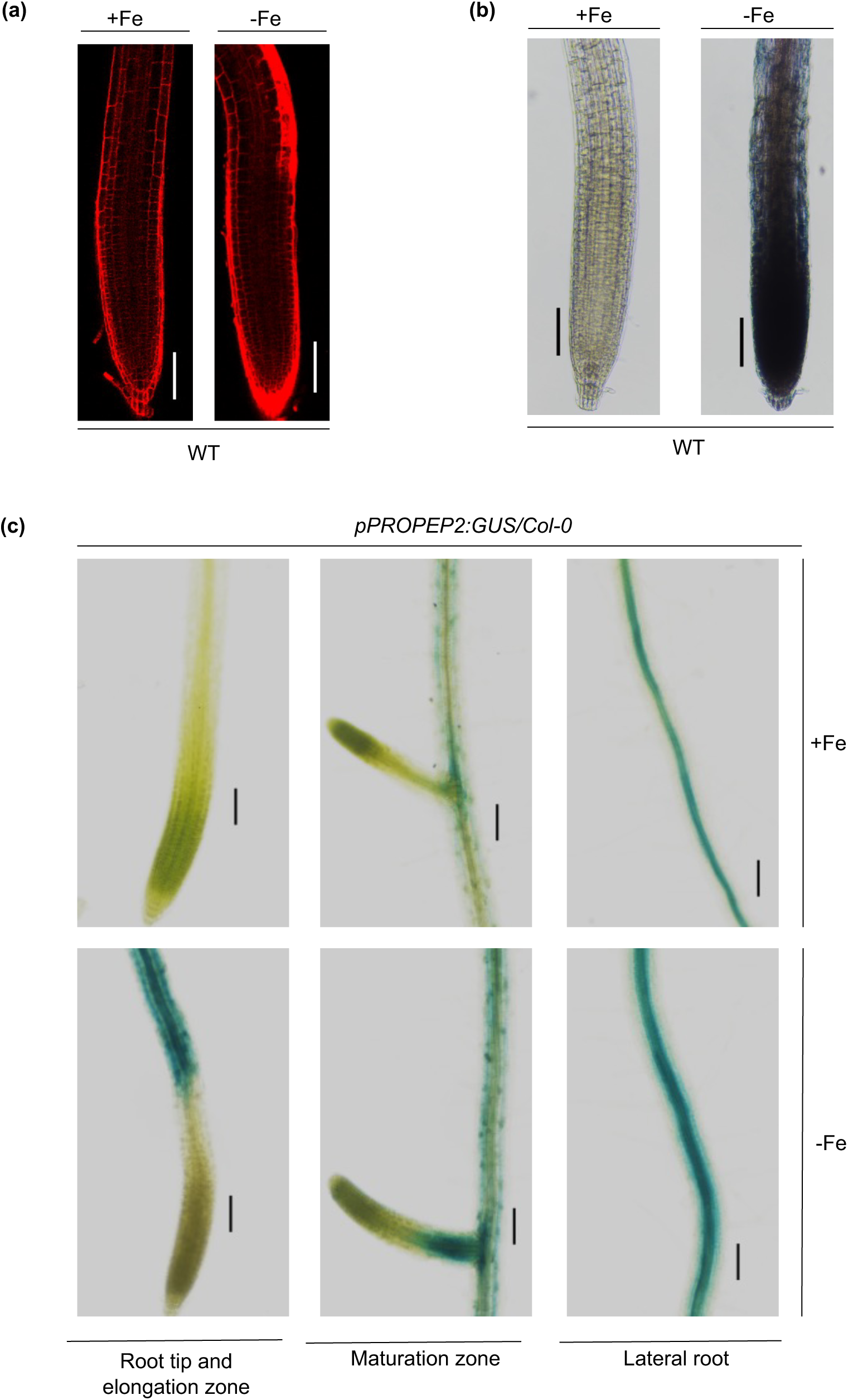
*PROPEP2* is induced under Fe deficiency. **(a-b)** Cell vitality is affected in roots of *A. thaliana* (*Col-0*) due to iron deficiency stress. Intense red and blue colors are indicative of cell death due to uptake of PI stain and Evans blue dye respectively. **(c)** Promoter activity of *PROPEP2* in roots of *pPROPEP2:GUS/Col-0* in response to +Fe and –Fe in lateral root, maturation zone, elongation zone and root tip regions. Five days old seedlings were transferred to +Fe and –Fe media for 72 hours and were subjected to confocal microscopy, Evans staining and GUS activity respectively. Scale bar: 100 µm

### Loss of function of PEP2 increases plant tolerance to Fe deficiency

Previous studies have shown that severe Fe-deficiency impairs the growth and architecture of WT plants, exhibiting drastic developmental effects such reduction in primary root length, a decrease in root and shoot biomass, and chlorophyll content (Gruber et al., 2013). To investigate the role of PEP2 in Fe homeostasis, we used a homozygous T-DNA insertion line for PEP2 and compared its phenotype with the WT/Col-0. Compared to WT, *pep2* had longer roots when grown vertically under Fe-deficient growth conditions (Figure 2a-b). Furthermore, we engineered transgenic lines overexpressing PEP2 precursor gene (PROPEP2) driven by the 35S promoter. These overexpressed PROPEP2 (*35S::PROPEP2 3.1* and *35S::PROPEP2 3.2*) displayed shorter root phenotypes under Fe deficient conditions similar to wild-type seedlings (Figure 2a-b). To examine root phenotype at the cell level, we compared the root meristem cortex cell numbers (MCN) between WT and *pep2* within a meristematic region starting from the quiescent center up to the cortex transition boundary (TB) after 72 hours of Fe deficiency. We found that in wild-type *Col-0* seedlings, root meristem cortex cell number (MCN) was significantly less in –Fe condition as compared to +Fe condition (Figure S2a-d). On the contrary, in *pep2* mutant no such reduction was observed. Furthermore, we looked at cell size in differentiation zone of roots of WT and *pep2* after 72 hours of Fe deficiency stress. We observed that cell size of differentiating cells in WT roots was highly impacted under Fe deficiency but in *pep2* mutant, the cell size in differentiation zone of Fe deprived roots was almost similar to that of plants grown in Fe sufficient media (Figure S2c). This led us to conclude that PEP2 negatively controls the meristem length and cell size in differentiation zone of root under iron deficiency.

**Figure 2.**
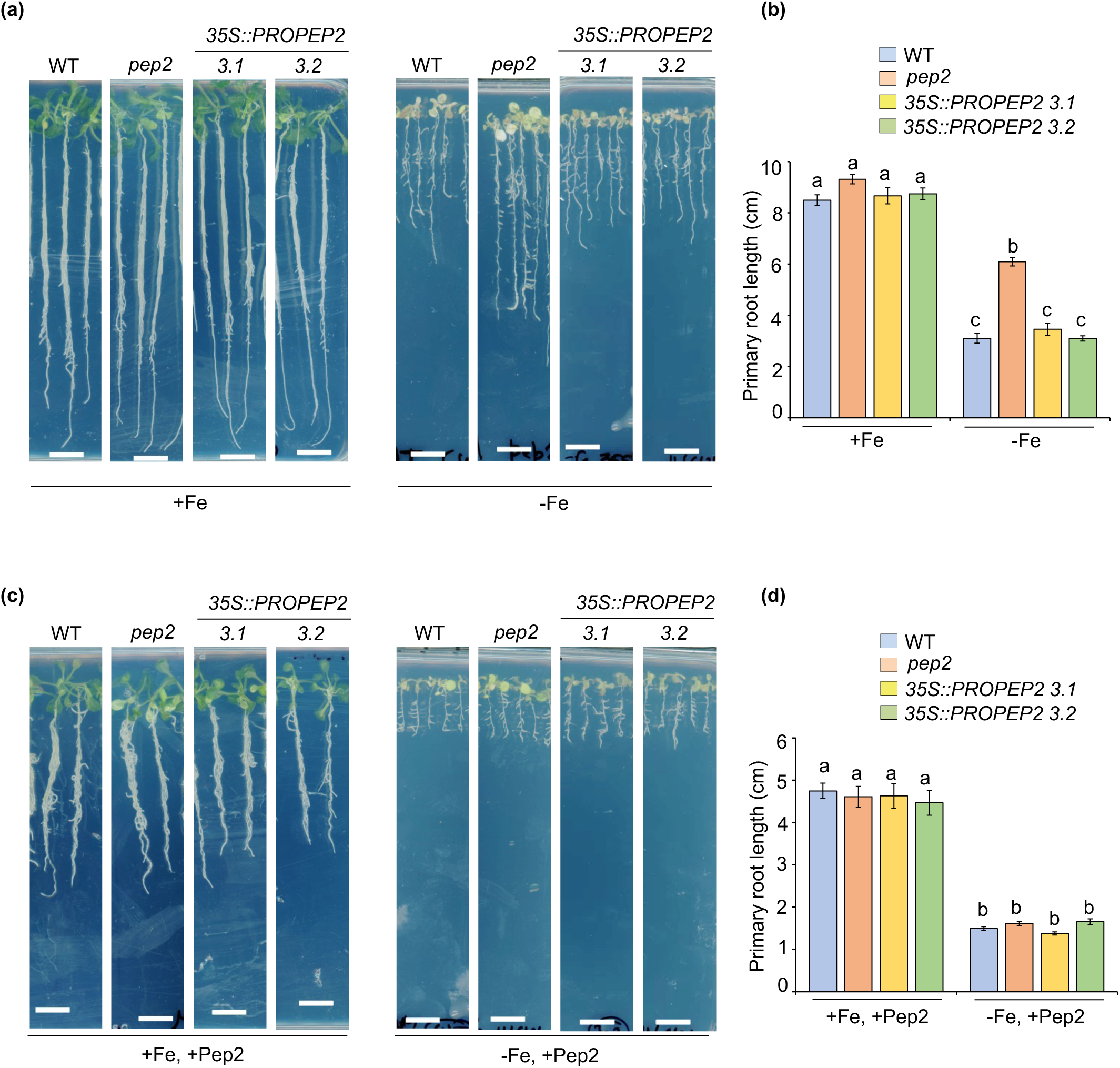
*PEP2* negatively regulates primary root growth under Fe deficiency. **(a-b)** Five days old seedlings of Arabidopsis wild-type (WT), *pep2* mutant and overexpression lines of PEP2 (*35S::PROPEP2 3.1* and *35S::PROPEP2 3.2)* transferred for nine days to liquid +Fe and -Fe media with 100µM Ferrozine. Scale bars: 1cm. **(b)** Bar graph of primary root length. **(c-d)** Five days old seedlings of Arabidopsis wild-type (WT), *pep2* mutant and overexpression lines of PEP2 (*35S::PROPEP2 3.1* and *35S::PROPEP2 3.2)* transferred for nine days to 50 nM Pep2 containing liquid +Fe and -Fe with 100µM Ferrozine. Scale bars: 1cm. **(d)** Bar graph of primary root length. Error bars represent average ± standard errors (SE). Different letters (a, b, c, and d) indicate significant differences, as determined by one-way ANOVA and a post hoc Tukey Test (P≤0.05).

Previous studies have demonstrated the role of Pep1 and Pep2 synthetic peptides in promoting root hair growth (Jing et al., 2024). Additionally, seedlings treated with synthetic Peptides cause a reduction in primary root growth (Jing et al., 2019, 2020). This led us to hypothesize that exogenous treatment of synthetic Pep2 might rescue the longer root phenotype of *pep2* under Fe deficiency. To prove this, we treated WT, *pep2* as well as *35S::PROPEP2* with 50 nM Pep2 under +Fe and –Fe conditions. We found that T-DNA mutant line as well as overexpression lines of PEP2 behaved similarly to the WT under both +Fe and –Fe conditions when supplemented with exogenous Pep2, showing similar primary root length. Thus, we conclude that the insensitive root phenotype of *pep2* mutant in response to –Fe stress can be rescued by exogenous application of the synthetic Pep2 (Figure 2c-d). Since, both severe –Fe stress and exogenous Peps negatively regulate root growth (Gruber et al., 2013; Jing et al., 2019, 2020), we attempted to explore the impact of synthetic peptide application on overall growth of the plant under –Fe condition. Our phenotypic analysis revealed a reduction in root and shoot fresh weight (RFW/SFW) as well as chlorophyll content in Fe starved WT seedlings as compared to +Fe condition (Figure S3a-d). Similar reduction is observed when WT seedlings were subjected to exogenous Pep2 under Fe sufficient condition and this reduction was more pronounced when Pep2 was applied to WT under –Fe stress condition (Figure S3a-d). These results suggest that PEP2 negatively regulates plant growth under Fe deficiency.

### PEP2 represses root response to –Fe by regulating the expression of Fe-deficiency-responsive genes

DAMP signalling and Fe deficiency trigger antagonistic responses in the root. For instance, Peps including Pep1 and Pep2 cause extracellular alkalinization which promotes plant immunity (L. Liu et al., 2022). However, Fe deficiency responses leads to an acidification of the rhizosphere to facilitate iron solubility (W. Schmidt et al., 2003). To investigate the role of PEP2 in regulating rhizosphere pH in response to –Fe, we screened Arabidopsis seedlings (WT, *pep2* and *35S::PROPEP2*) on medium with the pH indicator bromocresol. We found that under +Fe conditions, WT, *pep2* and *35S::PROPEP2* exhibited no acidified roots (Figure 3a-b). Meanwhile, iron deficiency triggered higher root acidification in *pep2* as compared to WT, whereas, *35S::PROPEP2* imparted root acidification similar to WT seedlings (Figure 3c-d). This suggested that the PEP2 loss of function leads to root acidification in response to -Fe.

**Figure 3.**
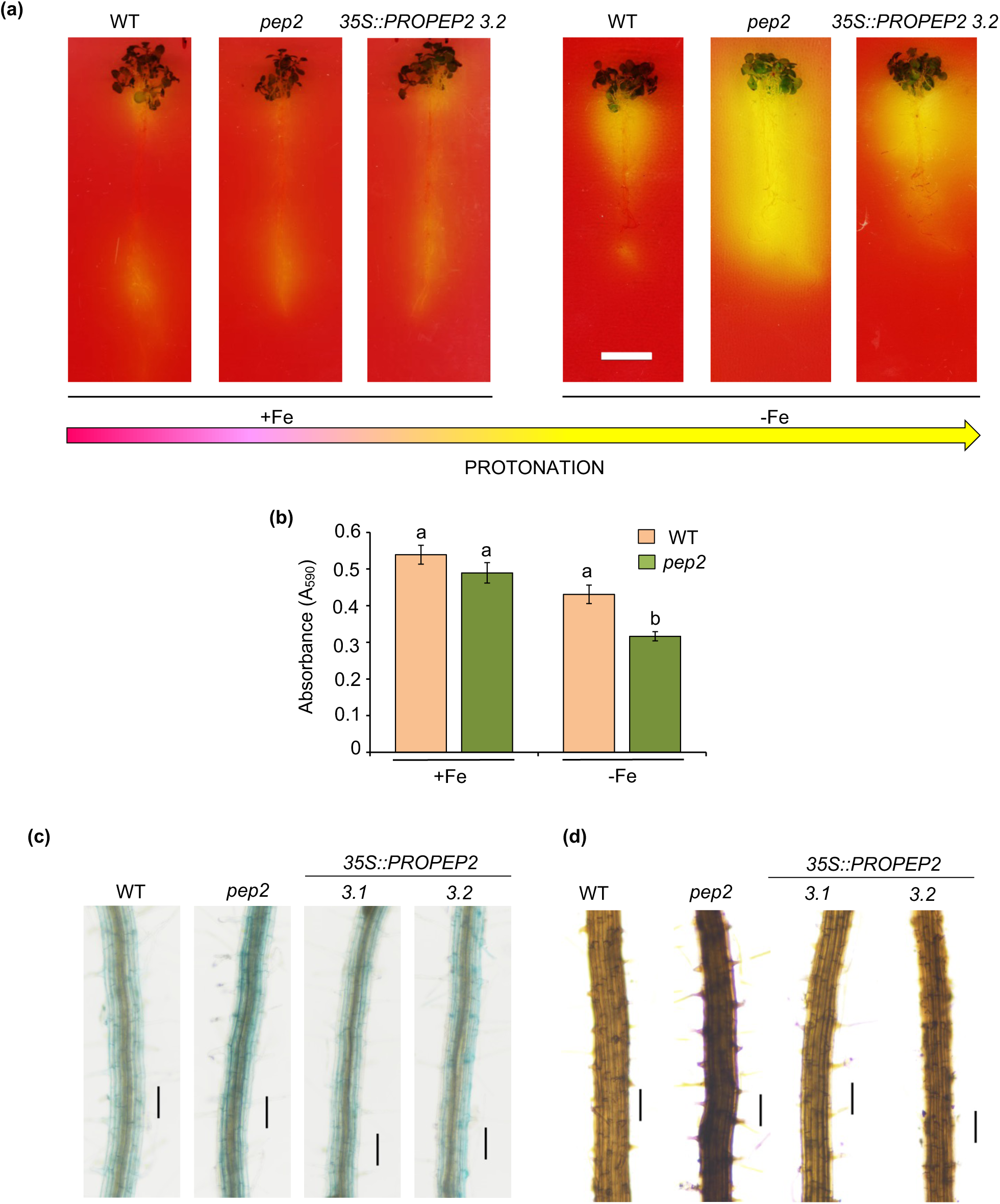
*PEP2* negatively regulates Fe accumulation and rhizosphere acidification. **(a)** Visualization of medium acidification in roots. Seven days old seedlings of WT, *pep2* and *35S::PROPEP2* were transferred to +Fe and –Fe media with 300µM Ferrozine for 72 hours and then transferred to rhizosphere acidification plates for 72 hours. Scale bar: 1cm. **(b)** Quantification of rhizosphere acidification responses of WT and *pep2* mutant plants. Bromocresol purple was used as the pH indicator. Error bars represent average ± standard errors (SE). Different letters (a and b) indicate significant differences, as determined by one-way ANOVA and a post hoc Tukey Test (P≤0.05). **(c)** Perls’ staining of maturation zone of five days old roots. **(d)** Perls’ DAB of maturation zone of five days old roots. Bar = 100µm.

To determine the role of PEP2 on Fe accumulation in vivo, we performed Perls (for Fe^3+^) and Perls/DAB (for Fe^2+^ and Fe^3+)^ staining. We found that *pep2* mutant roots accumulated more Fe as compared to WT. The transgenic line constitutively expression PROPEP2 under 35S (*35S::PROPEP2)*, also showed Fe content similar to WT but relatively less than *pep2* (Figure 3c-d). Therefore, we hypothesized that Pep2 abolishes root iron uptake. To further test this hypothesis, we measured ferric chelate reductase (FCR) activity and also an expression profile for some of the key Fe uptake genes including *IRT1* and *FRO2* under +Fe and –Fe conditions, with and without Pep2. Several studies have supported the molecular overlap between immunity and iron deficiency responses in plants (Herlihy et al., 2020; Yi Liu et al. 2021; Montejano-Ramírez & Valencia-Cantero, 2023; Romera et al., 2019; W. Schmidt et al., 2003). We observed an increase in expression levels of *IRT1* and *FRO2* in +Fe condition in response to Pep2 (Figure 4, 5), which was consistent with the previous studies as defense signals are known to modulate Fe uptake via inducing expression of Fe deficiency responsive genes including *IRT1*, *FRO2* and *PDR9* (Herlihy et al., 2020; Yi Liu et al., 2021; Montejano-Ramírez & Valencia-Cantero, 2023; Romera et al., 2019; W. Schmidt et al., 2003). While expression of *IRT1* as well as *FRO2* was significantly induced at 24 hours of Fe deficiency, treatment with Pep2 in –Fe condition abolished this induction (Figure 4a). Similarly, FCR activity in WT roots got induced under -Fe condition, and there was no induction observed in -Fe with Pep2 (Figure 4b). Likewise, lack of iron triggered the induction of the *pIRT1:4X:YFP* and *pIRT1:GUS/Col0* in the maturation zone of roots, and treatment with Pep2, abolished this response (Figure 4c-d, 5a). To further validate the role of PEP2 in regulating *IRT1* expression in planta, we crossed *pIRT1:GUS/Col0* with *pep2*. We compared *pIRT1:GUS* activity in the WT and *pep2* background grown on both +Fe and −Fe. We observed a stronger induction of the reporter gene in elongation zone, maturation zone as well as lateral roots of *pep2* as compared with the WT plants under −Fe conditions (Figure 5b). Collectively, these results indicate that under Fe deficiency, PEP2 negatively regulates the expression of *IRT1* and *FRO2* to repress iron uptake.

**Figure 4.**
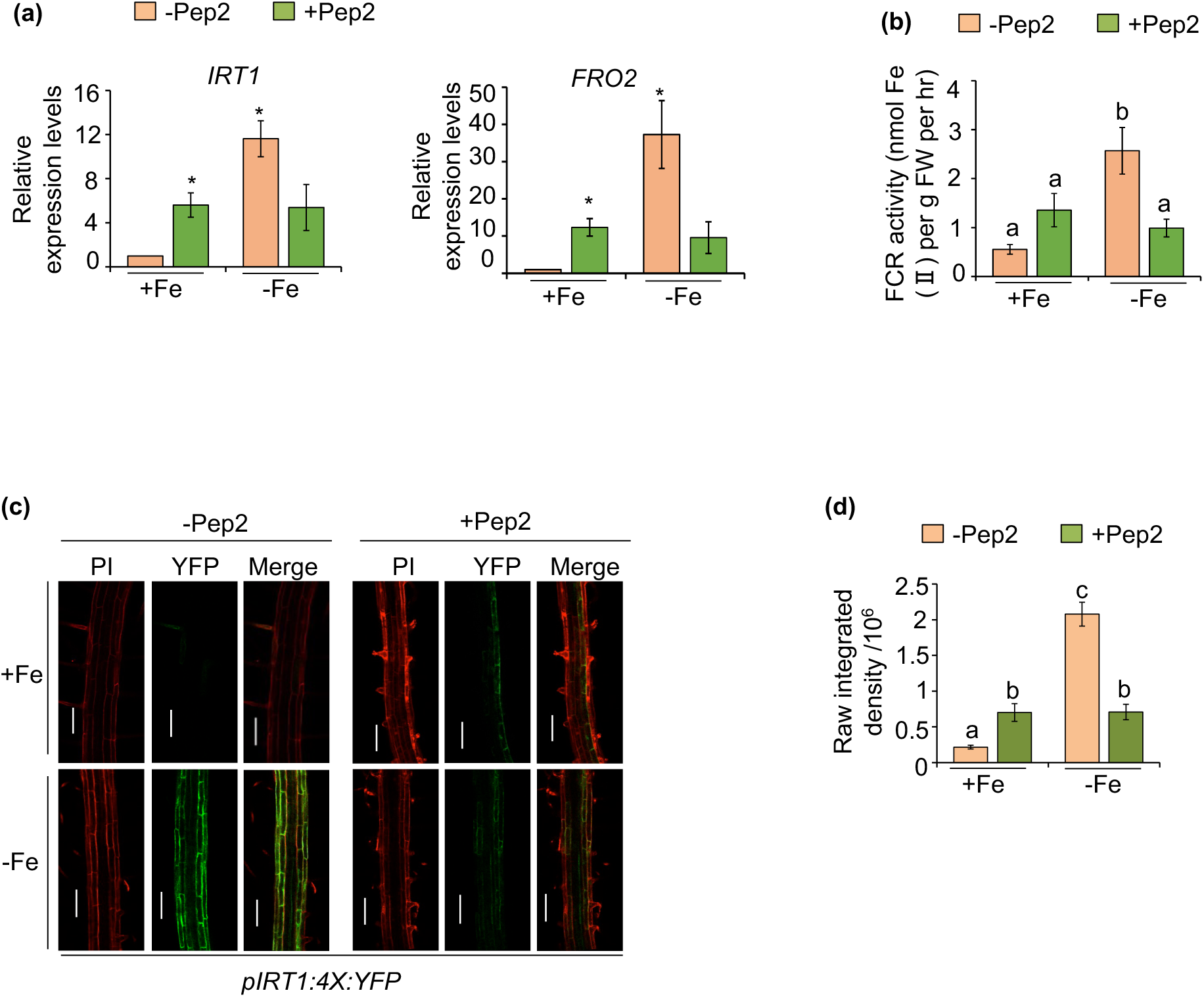
*PEP2* modulates expression of genes related to Fe homeostasis. **(a)** Expression levels of *IRT1* and *FRO2* in response to +Fe and –Fe with and without Pep2 by qRT-PCR. The gene expression level is normalized by TUB internal control. WT seedlings were treated with 1µM Pep2 for 24 hours. Data shown are an average of three biological replicates (n = 2 technical replicates). Each biological replicate consists of pooled RNA extracted from roots of ∼60 seedlings. Error bars represent ± SEM. Significant difference by Student’s t-test *(P ≤ 0.05) and **(P ≤ 0.01). **(b)** FCR activity in WT roots that were grown for six days on +Fe and then transferred for 72 hours to liquid +Fe and -Fe media with 300µM Ferrozine, with or without 100 nM Pep2. **(c)** Confocal images of *pIRT1:4X:YFP* and, **(d)** Quantification of Raw integrated Density of signal intensity in response to +Fe and –Fe, with and without Pep2. A representative single confocal section of maturation zone is shown. Scale bar: 100µm. Error bars represent average ± standard errors (SE). Different letters (a, b, c, and d) indicate significant differences, as determined by one-way ANOVA and a post hoc Tukey Test (P≤0.05).

**Figure 5.**
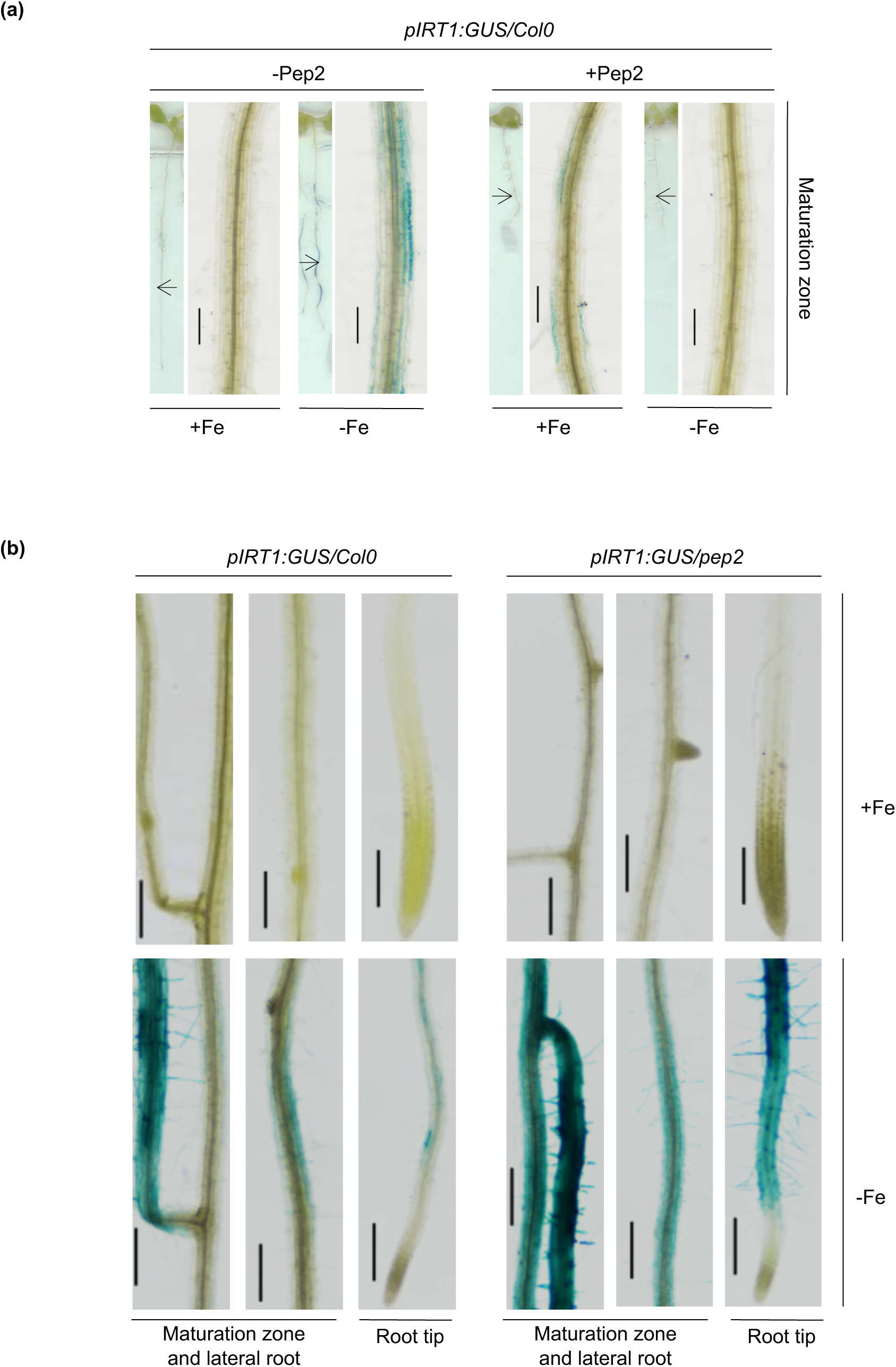
*PEP2* negatively regulates expression of *IRT1* under Fe deficiency. **(a)** Analysis of *pIRT1:GUS/Col0* expression in response to +Fe and –Fe with and without Pep2 treatment for 24 hours. Black arrows indicate magnified maturation zone regions. Scale bar: 100 µm. (b) Analysis of *pIRT1:GUS/Col0* and *pIRT1:GUS/pep2* expression in response to +Fe and –Fe after 72 hours. Scale bar: 100 µm.

Previous studies have shown that BRUTUS (BTS) and its two homologs (BTSL1 and BTSL2) negatively regulate iron uptake by suppressing the expression of several key genes, including *IRT1*, *AHA2* and *FRO2* (Y. Li et al., 2021; Selote et al., 2015). We hypothesized that BTS, BTSL1 and BTSL2 might mediate PEP2 triggered downregulation of key Fe uptake genes under its deficiency. To test whether these genes are involved in regulating Pep2-mediated Fe responses, we grew WT seedlings in +Fe and -Fe with and without Pep2 and checked the expression of *BTS*, *BTSL1* and *BTSL2*, which are known to get induced under –Fe conditions. We found that expression of *BTS* was significantly induced at 24 hours of Fe deficiency, and treatment of Pep2 enhanced this induction (Figure 6a). Also, no induction in the expression of *BTSL1* and *BTSL2* was observed in response to Pep2 under –Fe conditions (Figure 6b-c). This suggested that BTS and not BTSL1 and BTSL2 might regulate the Pep2-mediated repression of iron uptake. To further validate the role of PEP2 in regulating *BTS* expression, we analyzed the expression patterns of *pBTS:GUS/Col-0* and *pBTS:BTS:GFP* in +Fe and −Fe with and without Pep2 (Figure 5e,f). We observed a strong induction of the *BTS* in the stele of roots under Fe deficiency conditions, and Pep2 treatment further enhanced this response (Figure 6e,f). Also, the spatial expression patterns of BTS coincided with the area in which PEP2 is expressed (Figure 1c, 6e). Next, we wanted to check whether PEP2 controls Fe uptake through BTS. To test this, we checked the FCR activity in WT and *bts* mutant plants and found that as compared with WT, FCR activity in *bts* mutant remained insensitive to exogenous Pep2 under -Fe (Figure 6d). Taken together, our data indicates that BTS is likely to play a key role in regulating Pep2-induced suppression of the iron uptake. However, the mechanism behind this repression awaits further investigation.

**Figure 6.**
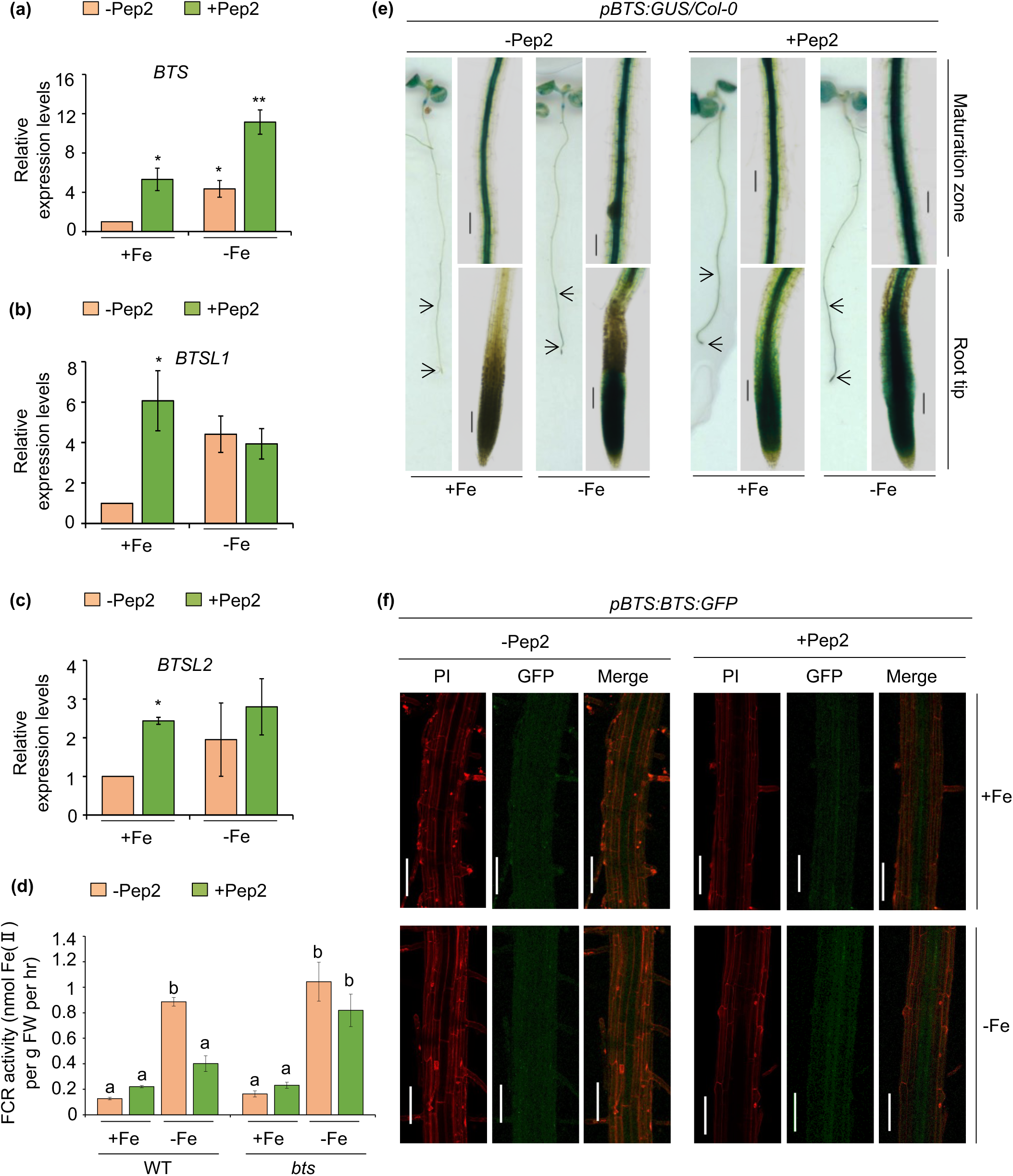
*PEP2* modulates expression of *BTS* under Fe deficiency. **(a-c)** Expression levels of BTS, *BTSL1* and *BTSL2* in response to +Fe and –Fe with and without Pep2 by qRT-PCR. The gene expression level is normalized by TUB internal control. The seedlings were treated with 1µM Pep2 for 24 hours. Data shown are an average of three biological replicates (n = 2 technical replicates). Each biological replicate consists of pooled RNA extracted from roots of ∼60 seedlings. Error bars represent ± SEM. Significant difference by Student’s t-test *(P ≤ 0.05) and **(P ≤ 0.01). **(d)** FCR activity in WT and *bts* roots that were grown for six days on +Fe media and then transferred for 72 hours on liquid +Fe and -Fe media with 300µM Ferrozine, with or without 100nM Pep2 **(e)** Analysis of *pBTS:GUS* gene activity in WT in response to +Fe and –Fe with and without Pep2 after 72 hours. Black arrows indicate magnified maturation zone and root tip regions. Scale bar: 100 µm. **(f)** Confocal images of *pBTS:BTS:GFP* in response to +Fe and –Fe with and without Pep2 after 72 hours. For each treatment, a representative single confocal section of maturation zone is shown. Scale bar: 100µm.

### *PEPR1* and *PEPR2* are induced under Fe deficiency

Peps are well documented components of the DAMP family that are recognized by a closely related receptor pair i.e PEPR1 and PEPR2, which are strongly induced by external abiotic stressors such as wounding and methyl jasmonate (MeJA) treatment (Bartels et al., 2013; Jing et al., 2019; Yamaguchi et al., 2010). A qRT-PCR was conducted using total RNA extracted from root of WT seedlings grown on +Fe and –Fe media to analyze *PEPR1* and *PEPR2* gene expression. Our qRT-PCR result showed a significant increase in *PEPR1* and *PEPR2* expression in response to −Fe conditions (Figure 7a). To examine their spatial expression patterns at the tissue level, we used transgenic lines with putative promoters fused to a β-glucuronidase (GUS) and green fluorescent protein (GFP) gene reporters respectively. GUS staining and confocal microscopy of *pPEPR1:GUS:GFP/Col-0* and *pPEPR2:GUS:GFP/Col-0* transgenic seedlings revealed extensive expression of both PEPRs in lateral root as well as maturation zone of root system under -Fe (Figure 7b-c). The expression in *pPEPR2:GUS:GFP/Col-0* seedlings were primarily localized to vascular tissue similar to *pPROPEP2:GUS/Col-0*, while the promoter of PEPR1 exhibited activity throughout the whole root (Figure 1b-c, 7c). Taken together, these results suggest, –Fe stress condition induces the transcription of *PEPR1* as well as *PEPR2* genes.

**Figure 7.**
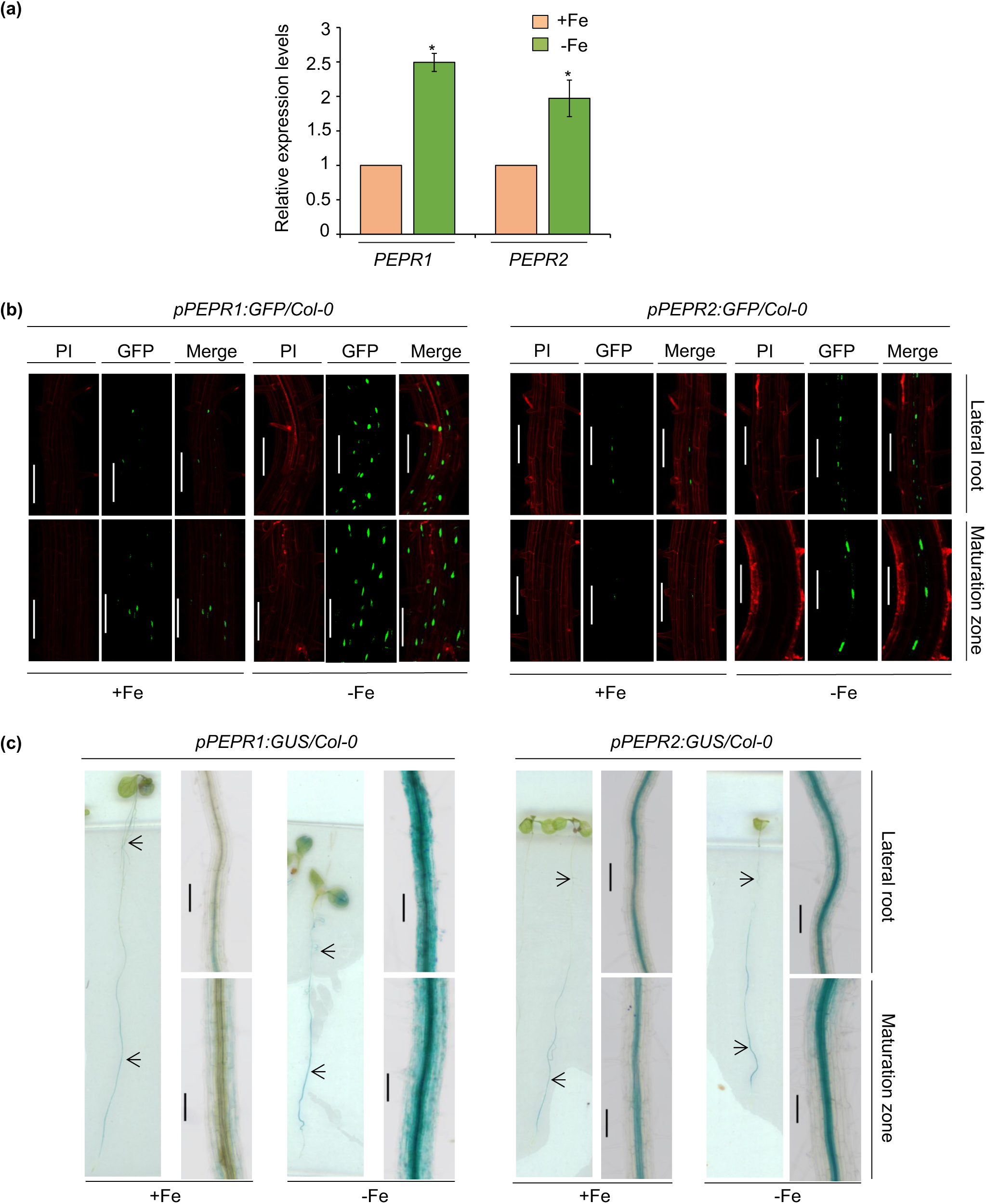
*PEPR1* and *PEPR2* are induced under Fe deficiency. **(a)** Expression levels of *PEPR1* and *PEPR2*. Relative expression was determined by qRT-PCR in wild-type (WT) grown on +Fe media for 6 days and transferred to +Fe and –Fe media with 300µM Ferrozine for 72 hours. The gene expression level is normalized by TUB internal control. Data shown are an average of three biological replicates (n = 2 technical replicates). Each biological replicate consists of pooled RNA extracted from roots of ∼90 seedlings. Error bars represent ± SEM. Significant difference by Student’s t-test *(P ≤ 0.05). **(b)** Promoter activity of *PEPR1* and *PEPR2* in roots of *pPEPR1:GFP/Col-0* and *pPEPR2:GFP/Col-0* seedlings respectively in response to +Fe and –Fe. Green: nuclear localized GFP signals; red: propidium iodide (PI) cell wall stain. In each treatment, the Z-stack scan is processed to maximal Z-projection (Merge, PI and GFP).Scale bar: 100µm. **(c)** Promoter activity of *PEPR1* and *PEPR2* in roots of *pPEPR1:GUS/Col-0* and *pPEPR2:GUS/Col-0* seedlings respectively in response to +Fe and – Fe. Black arrows indicate magnified maturation zone and root tip regions. Scale bar: 100 µm.

### PEP2-PEPR2 module negatively regulates primary root growth under -Fe conditions

Previous studies have highlighted the role of Pep-PEPR signalling in stimulating root hair development and inhibiting primary root growth (Jing et al., 2019, 2024; Zheng et al., 2018). As *PEPR1* and *PEPR2* are strongly induced in response to –Fe, to investigate their role in Fe homeostasis, we analyzed the root phenotype of the single mutant for PEPR1 (i.e. *pepr1*), PEPR2 (i.e. *pepr2*) and double mutant for PEPRs (i.e. *pepr1/2*) in response to low Fe. Compared to WT, all three mutant lines (i.e *pepr1*, *pepr2* and *pepr1/2*) displayed significantly longer roots when grown under −Fe as compared to the +Fe condition (Figure 8a-b). Our phenotypic analysis revealed that *pepr* mutants were less sensitive to −Fe as compared with the WT plants. A recent study on PROPEP2 has suggested that exogenous treatment of synthetic Pep2 results in a significant increase in both root hair density and length in WT and *pepr1* as compared to *pepr2*, with no stimulation of root hair growth in *pepr1/2* double mutant. This underscores the higher affinity of PEPR2 receptor to perceive exogenous Pep2 in controlling root hair growth (Jing et al., 2024). Further to check the potential receptor of Pep2 in regulating primary root length, we treated WT, *pepr1*, *pepr2* and *pepr1/2* with synthetic Pep2 and found that under both iron sufficient as well as iron deficient conditions, exogenous treatment of Pep2 inhibited primary root length in WT as well as *pepr1* mutant (Figure 8a-d). In contrast, neither *pepr2* nor *pepr1/2* showed any difference in primary root length when treated with Pep2 exogenously as compared with the WT and *pepr1* plants (Figure 8a-d). These findings indicate that PEPR2 perceives the Pep2 peptide and transduces this signal, leading to root growth inhibition and insensitive primary root length phenotype of *pepr2* in response to –Fe is due to impaired Pep2 perception via PEPR2 receptor.

**Figure 8.**
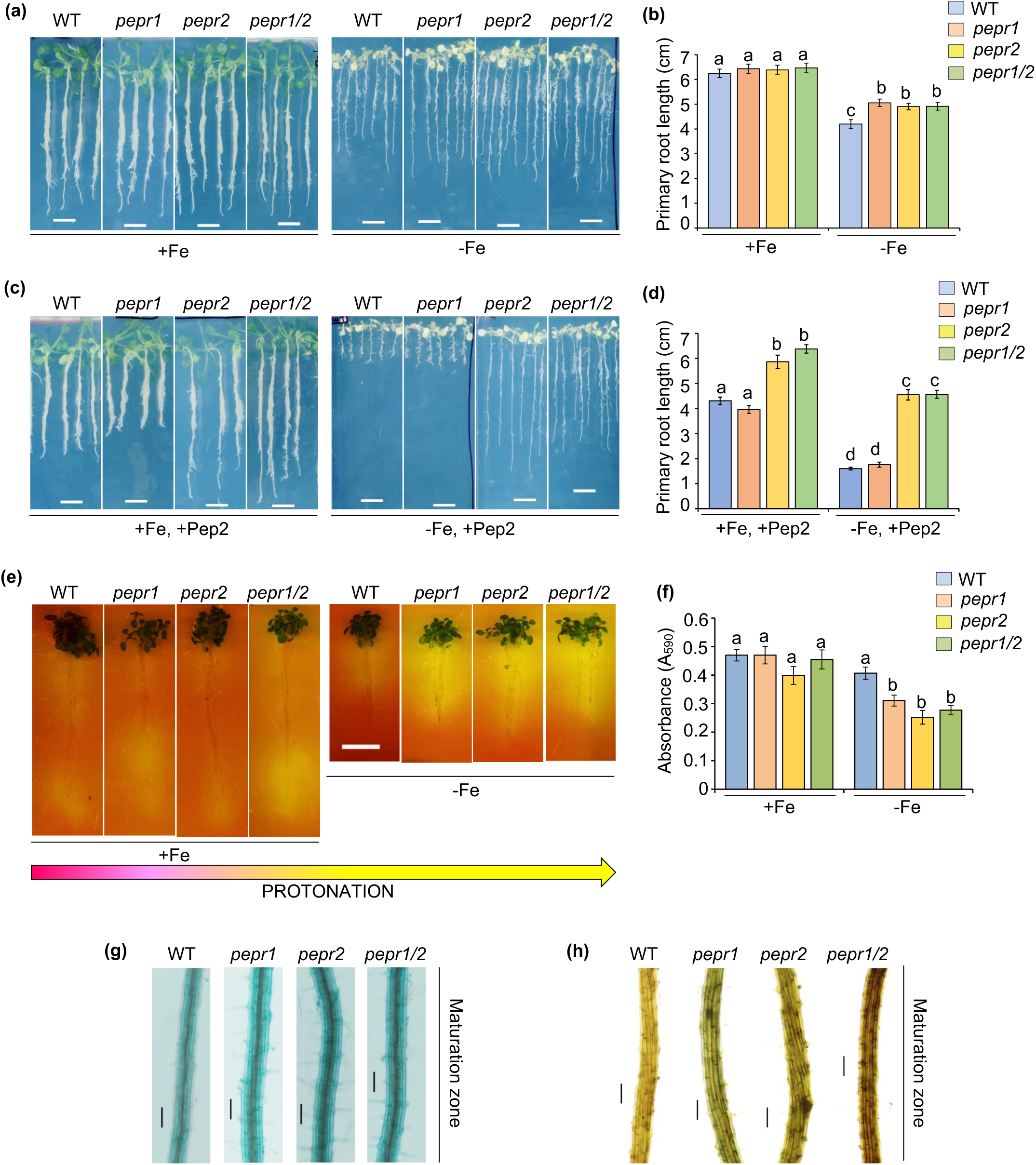
PEP2 is perceived by PEPR2 to negatively regulate primary root growth under Fe deficiency. **(a-b)** Five days old seedlings of Arabidopsis wild-type (WT) and single (i.e. *pepr1, pepr2*) as well as double (i.e. *pepr1/2*) mutants of PEPRs transferred for nine days to liquid +Fe and -Fe media with 100µM Ferrozine. Scale bars: 1cm. **(b)** Bar graph of primary root length. **(c-d)** Five days old seedlings transferred for nine days on Pep2 containing liquid +Fe and -Fe with 100µM Ferrozine. Scale bars: 1cm. **(d)** Bar graph of primary root length. **(e)** Visualization of medium acidification around roots. Seven days old seedlings were transferred to +Fe and –Fe media with 300µM Ferrozine for 72 hours and then transferred to rhizosphere acidification plates for 72 hours. Scale bar: 1cm. **(f)** Quantification of rhizosphere acidification responses of WT and *peprs*. Bromocresol purple was used as the pH indicator. **(g)** Perls’ staining of maturation zone of five days old roots. **(h)** Perls’ DAB of maturation zone of five days old roots. Bar = 100µm. Error bars represent average ± standard errors (SE). Different letters (a, b, c, and d) indicate significant differences, as determined by one-way ANOVA and a post hoc Tukey Test (P≤0.05).

Since, Fe deficient condition is known to trigger acidification of rhizosphere to enhance iron uptake (W. Schmidt et al., 2003), we therefore investigated the rhizosphere acidification capacity of *pepr1*, *pepr2* and *pepr1/2* mutants and compared it with WT under +Fe and –Fe conditions. We found that, under +Fe conditions, WT and *peprs* exhibited very little acidification, whereas significantly higher root acidification is found in *pepr* mutants as compared to WT under -Fe condition (Figure 8e-f). These data indicate that loss of function of PEPR1 and PEPR2 leads to increased root acidification that is initially triggered by -Fe. To determine the role of PEPR1 and PEPR2 on Fe accumulation in vivo, we performed Perls as well as Perls/Dab staining. We found that *pepr1*, *pepr2* and *pepr1/2* roots accumulated more Fe as compared with WT (Figure 8g-h). These results suggest that PEPRs negatively regulate rhizosphere acidification and iron accumulation under –Fe.

### PEP2-PEPR2 module regulates RBOHD mediated ROS accumulation under Fe deficiency

ROS acts as secondary messenger that influences plant stress responses against drought, salinity, and nutrient deficiency stress conditions (Jiao et al., 2013; Shin et al., 2005; Tsukagoshi, 2016). The PM-localized ROS generating enzyme i.e. respiratory burst oxidase homologs (RBOHs), are responsible for production of two types of ROS, including superoxide ion (O_2_^−^) and hydrogen peroxide (H_2_O_2_) (Sachdev et al., 2021). In Arabidopsis, out of ten members of RBOH family, RBOHD and RBOHF play a crucial role in apoplastic ROS generation in response to stress stimuli (Kadota et al., 2014; Torres et al., 2002). Several studies provide ample evidences for ROS mediated regulation of primary root length, root hair development and lateral root growth in plants (Shin et al., 2005; Tsukagoshi, 2016). In the previous studies, the Pep1-PEPRs signalling was shown to trigger ROS production via RBOHD activation (Jing et al., 2020; Kadota et al., 2014; Z. Liu et al., 2013). To evaluate whether roots mount a similar response on Pep2 peptide application, we first soak-incubated the WT and *pep2* seedlings in +Fe media with and without synthetic Pep2, followed by examining O_2_^−^ levels. By using the nitroblue tetrazolium (NBT) staining of O_2_^−^, we found that *pep2* mutants showed less level of O_2_^−^ accumulation than WT meanwhile Pep2 treatment strongly induced the accumulation of O_2_^−^ in the WT roots as compared to *pep2* roots (Figure 9a-b). However, such an increase was inhibited by potassium iodide (KI), a scavenger of ROS (Figure 9a-b). Role of ROS signalling is already known in –Fe (Reyt et al., 2015; Winterbourn, 1995). It was previously shown that knockout of stress responsive genes including *MNB1* and *ROP6* reduces ROS levels in response to –Fe (Song et al., 2022; Zhai et al., 2018). Thus, we hypothesized that the loss-of-function of PEP2 decreased the accumulation of ROS under Fe deficiency. To test this hypothesis, we used NBT staining to test changes in ROS levels in WT and *pep2* plants under control and Fe-deficiency conditions (Figure 9c-d). Under –Fe stress condition, a less intense staining of NBT was observed in *pep2* as compared to WT, suggesting lower levels of ROS accumulation in *pep2* mutant plants. Additionally, application of KI inhibited the increase in ROS accumulation in WT in response to –Fe (Figure 9c-d). The above results indicate that PEP2 promotes oxidative damage under –Fe stress conditions. Furthermore, to evaluate whether Pep2 induced ROS accumulation is via PEPR1 or PEPR2, we treated WT, *pepr1*, *pepr2*, *pepr1/2* and *pep2pepr2* double mutant with synthetic Pep2 under iron sufficient condition and detected the ROS levels in the roots. By using the nitroblue tetrazolium (NBT) staining of O_2_^−^, we found that Pep2 treatment effectively enhanced the production of O_2_^−^ in WT and *pepr1* mutants. In contrast, *pepr2*, *pepr1/2* and *pep2pepr2* mutants were insensitive to Pep2 triggered ROS accumulation (Figure 9e-f). These results suggest that Pep2-PEPR2 module functions in roots to mediate ROS production and regulate root growth under iron deficiency.

**Figure 9.**
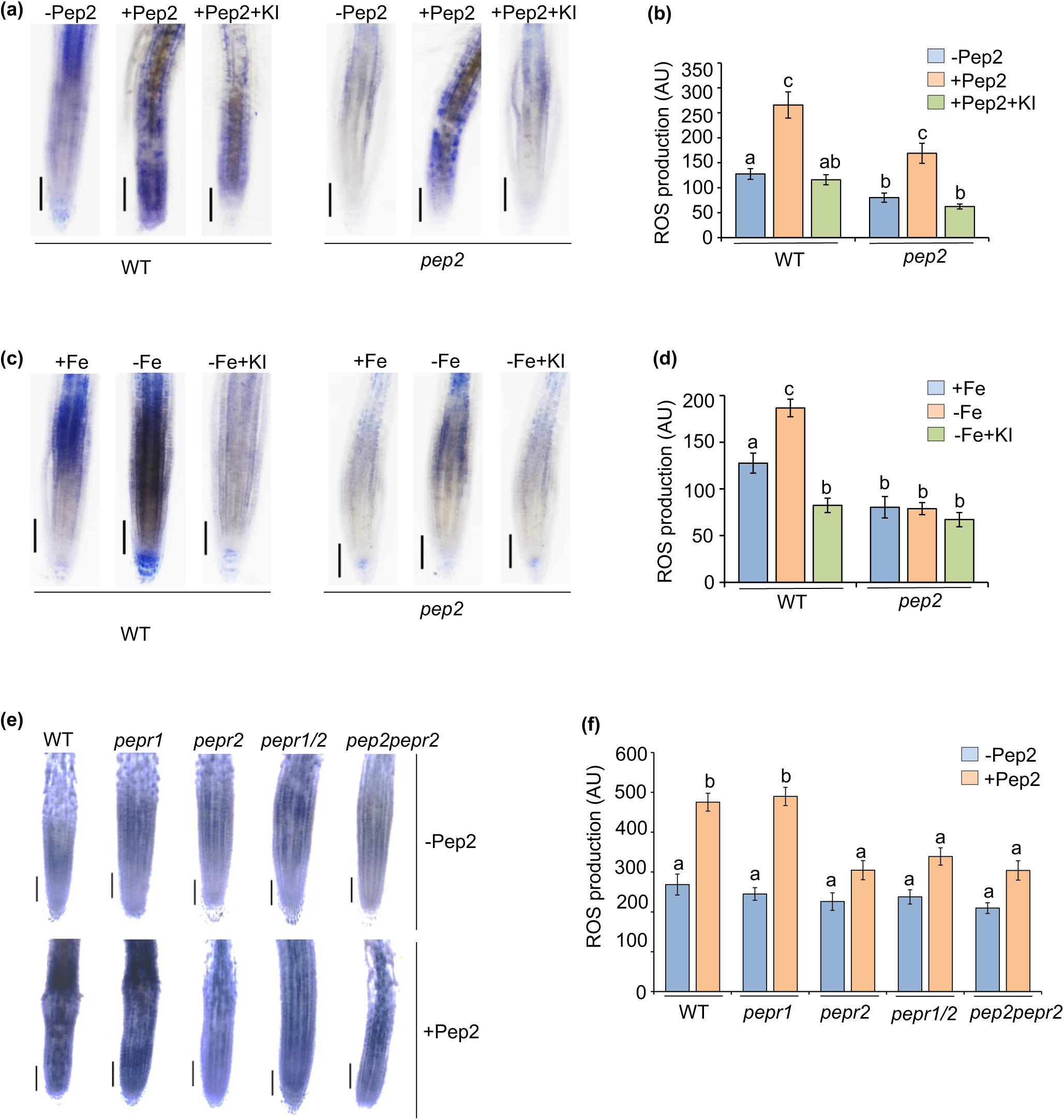
PEP2-PEPR2 module affects ROS accumulation under Fe deficiency. **(a,b)** Five days old seedlings of WT and *pep2* mutants were transferred to liquid media i.e. +Fe, +Fe+ 100nM Pep2 and +Fe+ 100nM Pep2+ 200µM KI for 24 hours and subjected to NBT staining for superoxide. Scale bar: 100µm. **(b)** Bar graph for ROS production in root tip. AU (Arbitrary unit). **(c-d)** Five days old seedlings of WT and *pep2* mutants were transferred to liquid +Fe, -Fe media with 100µM Ferrozine and -Fe with 100µM Ferrozine+ 200µM KI for 24 hours and subjected to NBT staining. **(d)** Bar graph for ROS production in root tip. **(e-f)** Five days old seedlings of WT, *pepr1*, *pepr2*, *pepr1/2* and *pep2pepr2* mutants were transferred to liquid +Fe and +Fe+ 1µM Pep2 for 24 hours and subjected to NBT staining. Scale bar: 100µm **(f)** Bar graph for ROS production in root tip. Error bars represent average ± standard errors (SE). Different letters (a, b, c, and d) indicate significant differences, as determined by one-way ANOVA and a post hoc Tukey Test (P≤0.05).

Furthermore, to test the function of RBOHs in Pep2-PEPR2 signalling under iron deficiency, we first analyzed the transcriptional levels of *RBOHD* and *RBOHF* in response to +Fe and –Fe conditions with and without the elicitor Pep2. Based on our qRT-PCR results, we concluded that expression of *RBOHD* was more induced than *RBOHF* on Pep2 elicitation under +Fe; moreover, this induction in *RBOHD* expression is even more pronounced when Pep2 is supplemented in Fe deficient media (Figure 10a-b). Since, the strong induction of *RBOHD* by the Pep2 is concomitant with ROS production, to prove this, we treated the WT, *rbohd* and *rbohf* seedlings with Pep2 in the presence as well as absence of a ROS scavenger, KI. We examined O_2_^−^ accumulation on Pep2 treatment using NBT staining, and found that *rbohd* mutant roots accumulated less level of O_2_^−^ than WT and *rbohf*, whereas KI scavenged Pep2 triggered O_2_^−^ accumulation in WT as well as *rbohf* mutant roots (Figure 10c-d). We further tested the accumulation of ROS in *rboh* mutants in response to –Fe stress condition and found a significant lower level of ROS accumulation in *rbohd* mutant plants when compared with WT and *rbohf* plants (Figure 10e-f). Collectively, these results suggest that PEP2-PEPR2 module promotes RBOHD-mediated ROS accumulation under Fe deficiency.

**Figure 10.**
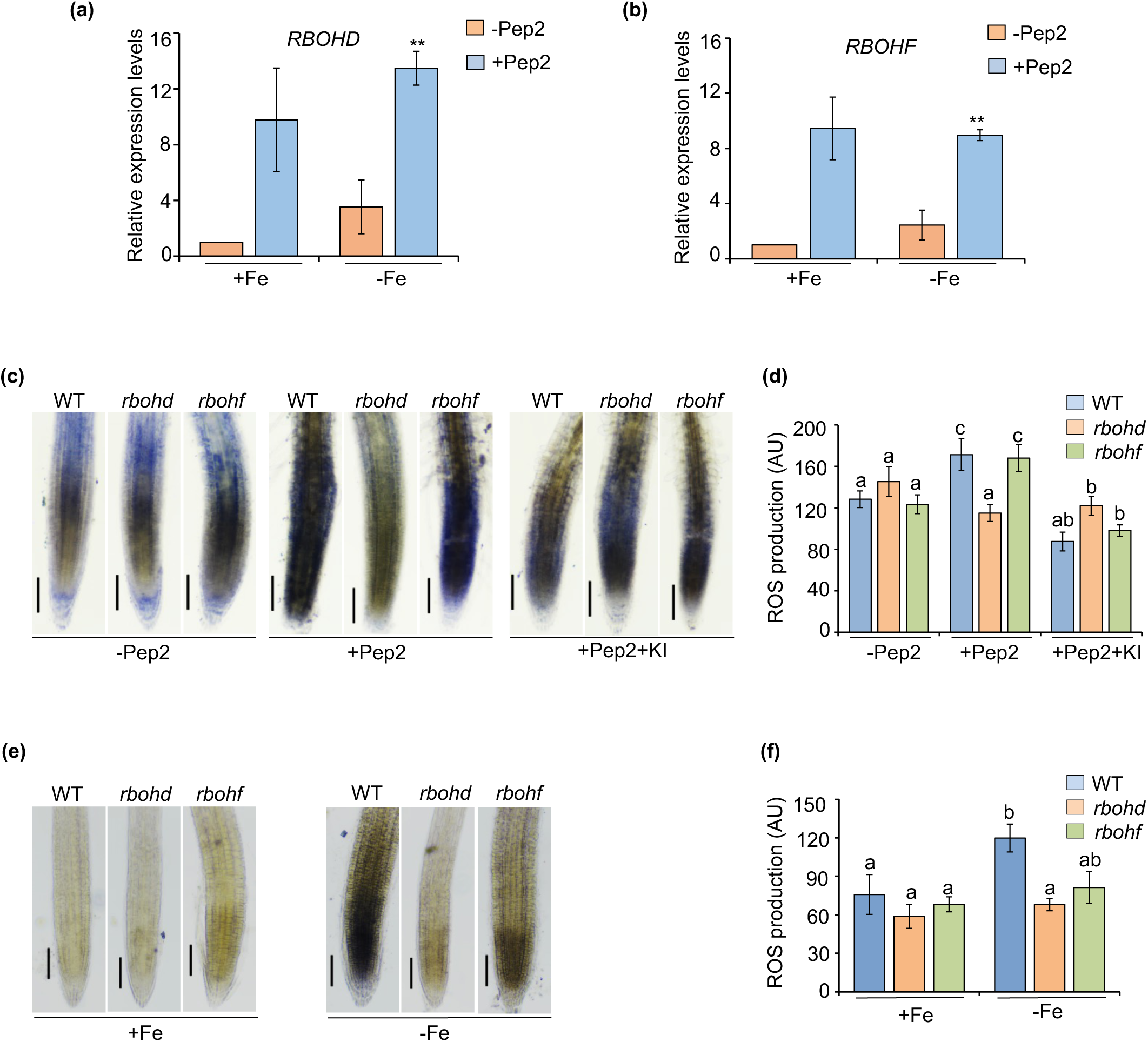
*PEP2* induced ROS accumulation is *RBOHD* mediated. **(a,b)** Expression analysis of *RBOHD* and *RBOHF* in response to +Fe and –Fe with and without Pep2 by qRT-PCR. The gene expression level is normalized by TUB internal control. The seedlings were treated with 1µM Pep2 for 24 hours. Data shown are an average of three biological replicates (n = 2 technical replicates). Each biological replicate consists of pooled RNA extracted from roots of ∼60 seedlings. Error bars represent ± SEM. Significant difference by Student’s t-test *(P ≤ 0.05) and **(P ≤ 0.01). **(c-d)** Five days old seedlings of WT and *rboh* mutants were transferred to liquid media i.e. +Fe, +Fe+ 1µM Pep2 and +Fe+ 100nM Pep2+ 200µM KI for 24 hours and subjected to NBT staining. Scale bar: 100µm. **(d)** Bar graph for ROS production in root tip. AU (Arbitrary unit). **(e-f)** Five days old seedlings of WT and *rboh* mutants were transferred to solid +Fe and -Fe media for 24 hours and subjected to NBT staining. **(f)** Bar graph for ROS production in root tip. Error bars represent average ± standard errors (SE). Different letters (a, b, c, and d) indicate significant differences, as determined by one-way ANOVA and a post hoc Tukey Test (P≤0.05).

## DISCUSSION

Fe is an essential mineral element and its deficiency leads to detrimental effects on plant growth resulting in reduced crop yield. To overcome adverse effects, plants have developed complicated regulatory networks to modulate Fe homeostasis and deficiency responses. Many signalling molecules including plant hormones affect Fe deficiency response in both graminaceous and non-graminaceous plants, though the regulatory function of signalling peptides and their receptors is less explored (Abadía et al., 2011; Kobayashi & Nishizawa, 2012b; Marschner & Römheld, 1994). A recent study has shown that flg22 MAMP participates during Fe deficiency response establishment in Arabidopsis (Cao et al., 2024). To date, it is not clear how plant system adapts to low Fe availability in response to DAMP signals. Our work provides insights into the molecular overlap between DAMP and Fe signalling and uncovers a molecular player of Fe signalling, the PEP2 endogenous peptide (Figure 11).

**Figure 11.**
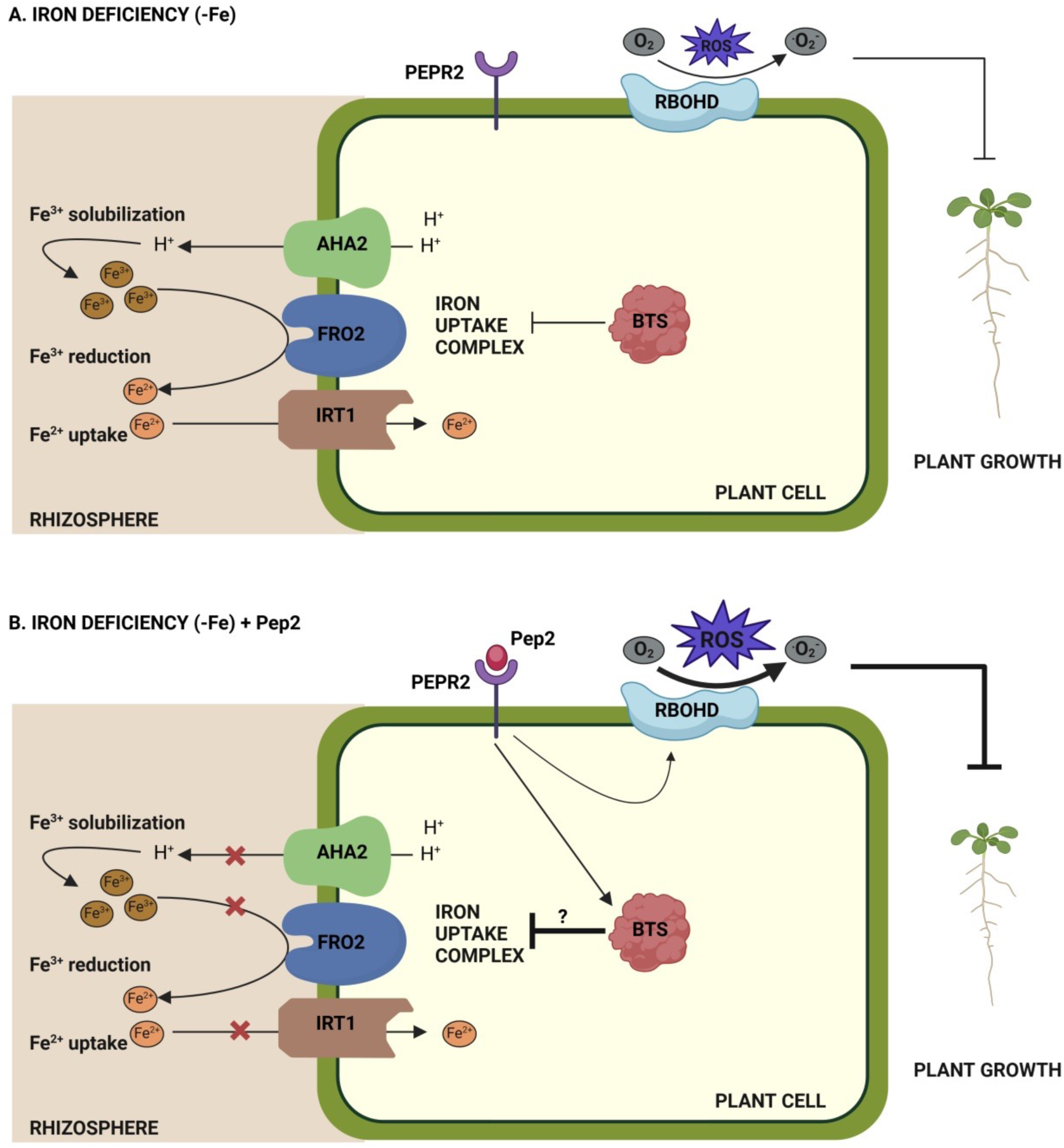
Working model of the role of PEP2-PEPR2 module in regulating Fe homeostasis and plant growth under Fe deficiency in *Arabidopsis thaliana*. **A.** Under Fe deficiency stress (-Fe), in the absence of Pep2 perception via PEPR2 receptor, iron uptake complex comprising of *AHA2*, *FRO2* and *IRT1* is activated. *AHA2* increases the acidification of rhizosphere to facilitate Fe^3+^ solubilization. Solubilized Fe^3+^ is then reduced from Fe^3+^ to Fe^2+^ by *FRO2* which is then imported into the root cell via *IRT1*. *BTS* expression is also induced in response to -Fe which limits the activity of iron uptake complex. Moreover, ROS accumulation is induced via *RBOHD* in response to –Fe, which leads to plant growth inhibition. **B.** Under Fe deficiency, in response to Pep2 binding to PEPR2 receptor, iron uptake complex is inactivated while *BTS* expression is strongly induced. This increase in *BTS* expression further represses the activity of iron uptake complex. However, the mechanism behind this repression awaits further investigation. Pep2-PEPR2 signaling has an additive effect on –Fe induced ROS accumulation which further limits plant growth. Positive and negative regulatory actions are indicated by arrows and lines with bars, respectively. Arrow thickness represents the amount of product formed. Figure created with biorender.com. Abbreviations: Fe, Iron; Pep2, Plant Elicitor Peptide 2; PEPR2, Plant Elicitor Peptide Receptor 2; Fe^3+^, Ferric; Fe^2+^, Ferrous; AHA2, Autoinhibited-H^+^-ATPases 2; FRO2, Ferric-Reduction Oxidase 2; IRT1, Iron Regulated Transporter 1; BTS, BRUTUS; ROS, Reactive Oxygen Species; RBOHD, Respiratory Burst Oxidase Homologue D.

Previous research has primarily focused on the effect of Pep2 in shaping overall root system architecture, which has contributed significantly to enrich our understanding of the functions of PEP2 in modulating plant defense responses (Jing et al., 2024). In the present research, we demonstrate that Arabidopsis plant elicitor peptide PEP2 negatively regulates Fe accumulation and plant tolerance in response to Fe deficiency. Since Peps are known to get released in response to cell damage via wounding and herbivory attack, we thereby checked cell vitality of WT roots in response to Fe deficiency. We found that cell vitality do get affected under –Fe, which might trigger the release of Pep2 as stress response (Figure 1). Furthermore we cross-confirmed this at tissue level and found a significant increase in PEP2 expression in root vasculature in response Fe starvation (Figure 1). Additionally, our phenotypic data suggests that knockdown of PEP2 displayed enhanced tolerance to –Fe with increased primary root growth, whereas *35S::PROPEP2* plants showed severe sensitivity to –Fe just like WT/*Col-0* seedlings (Figure 2a-b). Meanwhile, exogenous treatment of Pep2 increased *pep2* sensitivity to –Fe stress (Figure 2c-d). Our findings also indicated significantly higher root acidification in *pep2* plants as compared to WT and *35S::PROPEP2* plants under –Fe conditions (Figure 3a-b). As an acidic condition generally promotes Fe solubility in rhizosphere, our Perls and Perls/Dab staining results revealed that *pep2* accumulates more Fe as compared to WT and *35S::PROPEP2* (Figure 3c-d). Collectively, these results suggest that PEP2 negatively regulates rhizosphere acidification and Fe accumulation in roots to inhibit plant growth under Fe deficiency.

According to earlier studies, Fe-deficient stress responses are governed by a complex regulatory network of Fe uptake genes including *IRT1* and *FRO2* (Eide et al., 1996; Robinson et al., 1999; Santi & Schmidt, 2009; Stringlis et al., 2019). Here, we found that expression of these genes is induced under -Fe conditions and Pep2 application suppresses this induction (Figure 4,5). On contrary, Pep2 application promoted the induction of *BTS*, a negative regulator of Fe uptake, in response to low Fe stress (Figure 6). Additionally, *bts* mutants were insensitive to reduction in FCR activity in response to Pep2 as compared with WT (Figure 6). Taken together, these results indicate that Pep2 might negatively regulates Fe uptake by regulating BTS expression levels. However, further studies are required to explore this regulatory mechanism.

In the present work, we also investigated whether PEPR1 and PEPR2 play a role in maintaining Fe homeostasis. Our phenotypic results suggest that single as well as double mutants of PEPRs were insensitive to –Fe conditions, showing longer primary root length, higher rhizosphere acidification and higher Fe accumulation than WT (Figure 8). Importantly, Pep2 is primarily perceived by PEPR2 to negatively regulate root growth in response to low Fe (Figure 8).

It is well known that ROS accumulation is induced by Pep signalling and –Fe stress (Jiao et al., 2013; Shin et al., 2005; Song et al., 2022; Tsukagoshi, 2016; Zhai et al., 2018). In our results, an exogenous supply of Pep2 also led to an enhanced accumulation of ROS in WT and *pepr1* roots than *pepr2* and *pep2pepr2* roots (Figure 9). Meanwhile *pep2* mutant plants were less sensitive to reactive oxygen damage than WT in response to exogenous Pep2 treatment and –Fe stress conditions (Figure 9). Since, NADPH oxidase including RBOHD and RBOHF, is known to generate ROS in response to stress conditions and Pep application (Jing et al., 2020; Kadota et al., 2014; Z. Liu et al., 2013). Here, we have confirmed a higher *RBOHD* gene expression in Pep2 treated WT roots as compared to *RBOHF* expression under –Fe condition (Figure 10). Moreover, NBT staining results suggest that *rbohd* roots accumulated lower level of (superoxide) O_2_^−^ than WT and *rbohf* in response to both Pep2 treatment and – Fe stress conditions (Figure 10). Taken together, these results suggest that Pep2-PEPR2 module functions in roots to promote RBOHD-mediated ROS accumulation and thereby inhibit root growth in response to Fe deficiency (Figure 11).

Based on our results and existing literature, we propose a model depicting the crucial involvement of PEP2 in modulating Fe-deficiency responses in Arabidopsis (Figure 11). PEP2-PEPR2 signalling module negatively regulates the expression of Fe uptake complex (*IRT1*, *AHA2* and *FRO2*), and positively regulates the expression of *BTS* under –Fe conditions (Figure 11). Under Fe deficiency, upon Pep2 binding to PEPR2 receptor, –Fe induced ROS accumulation via RBOHD becomes more pronounced which further limits plant growth (Figure 11). In summary, our findings provide novel evidence that PEP2 regulation of Fe-responsive genes through ROS generation is critical for modulating Fe-deficiency responses in Arabidopsis. Further experiments will be required to understand whether PEP2-PEPR2 module interacts with bHLH TFs involved in the Fe-signalling pathway. Since, PEP-PEPR signalling is known to mount immune responses in plants, in future, it will be interesting to explore the extent of cross link between plant stress responses and nutritional immunity. In light of this, unraveling the signalling pathways downstream of PEP-PEPR crosstalk becomes crucial for a holistic understanding of Fe homeostasis. Furthermore, the role of other candidate PEPs and identification of novel receptors or co-receptors will provide valuable insight into the mechanism by which Peps regulate root growth in response to –Fe or other nutrient deficiencies in Arabidopsis. Furthermore, the knowledge gained from Arabidopsis model plant can be translated to tomato, wheat, maize and other crop plants, as most of the stress-signalling pathways are evolutionally conserved.

## Materials and methods

### Plant materials and growth conditions

For all experiments, *Arabidopsis thaliana* ecotype Columbia (*Col-0*) was used as wild-type (WT) control. The mutant/transgenic lines used in this study were *pepr1*, *pepr2*, *pepr1/2*, *pPEPR1:GUS:GFP/Col-0* and *pPEPR2:GUS:GFP/Col-0* (Nakaminami et al., 2018), *pep2* (SALK_ 206498C), *rbohd* and *rbohf* (Kadota et al., 2014), *bts* (Long et al., 2010), *pIRT1:4X:YFP* (Martín-Barranco et al., 2020), *pIRT1:GUS/Col0* (Blum et al., 2014), *pBTS:BTS:GFP* and *pBTS:GUS* (Selote et al., 2015). The *pep2pepr2* and *pIRT1:GUS/pep2* lines were generated by crossing. The single and double-mutants were confirmed by genotyping. The primers used for genotyping are listed in Table S1. Seeds were surface sterilized, stratified in dark for 4 days and grown vertically on half-strength Murashige and Skoog (MS) medium including Fe salts (Caisson Labs, Smithfield, UT, USA) with final concentration of 1% sucrose and 1% agar as standard growth medium and half-strength Murashige and Skoog (MS) medium excluding Fe salts (Caisson Labs). The pH of media was adjusted to 5.75 using KOH. [3-(2-pyridyl)-5,6-diphenyl-1,2,4-triazine sulfonate] FerroZine (FZ) (Sigma-Aldrich, Bangalore, India) was added to create Fe-deficient conditions. For seedling growth, plants were grown under long-day conditions at 22^°^C, 60% humidity under 16h/8h light/dark condition with a light intensity of 90-110 µmolm^−2^s^−1^. Soil was made as a mixture of soilrite, perlite and compost in 3:1:1/2 composition.

### Plasmid constructs and transgenic lines

To generate overexpression line of *PROPEP2*, the coding sequence was amplified from cDNA. Obtained fragment was introduced into pENTR/D-TOPO vector using gateway cloning. The resulting clone was sequence verified and combined with destination vector (*pMDC32*) using the gateway LR reaction. The construct was introduced into *Agrobacterium tumifaciens* GV3101 which was finally used to transform WT/*Col-0* plants via floral-dipping. The transformed WT plants were selected via Hygromycin resistance to obtain homozygous T3 transgenic lines.

For *pPROPEP2:GUS/Col-0* construct, the promoter sequence of *PROPEP2* gene was amplified from genomic DNA and cloned in pENTR/D-TOPO. The resulting clone was used to set up a LR reaction with PKGWFS7 to create *pPROPEP2:GUS/Col-0*, which was then transformed into WT and T3 transgenic lines were selected on Kanamycin. The primers used are listed in Table S1.

### Elicitor preparation and treatment

Pep2 oligopeptide (DNKAKSKKRDKEKPSSGRPGQTNSVPNAAIQVYKEDWT) was synthesized by the S BioChem India. The Pep2 synthetic peptide was dissolved in deionized water. For elicitor treatment, Pep2 was added to the liquid media to obtain the respective final concentration (depending on the time scale of the assay 1 µM, 50 nM or 100 nM as described in the figure legends).

### Phenotypic analyses

For phenotypic characterization, 5 days old seedlings were transferred from +Fe (MS with agar) to liquid medium containing +Fe (MS) and –Fe (100µM FZ) respectively for 8 days with or without 50 nM Pep2 in 12 wells culture plates. The plates were kept at mild shaking in the growth chamber, and seedlings were scanned after 8 days of transfer using Epson Perfection V600 at 1200□dpi resolution. The root length was quantified using ImageJ 1.52a software (National Institutes of Health).

### Ferric chelate reductase (FCR) assay

The FCR assay was performed as previously described by Yi and group with some modifications (Yi & Guerinot, 1996). 6-day-old seedlings grown on +Fe (MS) plates were transferred to liquid medium with +Fe (MS), and –Fe (300µM FZ), with or without 100 nM Pep2 and treated for 72 hours. The seedlings were washed three times with water and roots of 10 seedlings were pooled as one sample. The samples were incubated in 700□μl of FCR assay solution (0.1□mM Fe (III)-EDTA and 0.3□mM ferrozine) and incubated for 1 hour in dark. The FCR activity was measured by spectrophotometer with the absorbance at 562 nm. The results were calculated using the formula [nmol Fe(□) per g FW per hr□=□(A/28.6)□×□V/Root FW].

### Total chlorophyll content measurement

The chlorophyll content was measured using previously described protocol with some modifications (Foyer et al., 1995). 5-day-old seedlings grown on +Fe (MS) plates were transferred to liquid medium with +Fe (MS), and –Fe (100µM FZ), with or without 50 nM Pep2 and treated for 8 days. Two shoots of plants were harvested and pooled as one sample. Chlorophyll was extracted using 1□ml of 80% acetone from leaf tissues and incubated in dark for 24□hours. Total chlorophyll content was calculated using the formula: (mg/g FW)□=□(20.3□×□A645□±□8.04□×□A663)□×□V/W□×□103.

### Confocal microscopy with PI staining

For cell type phenotype of WT and *pep2,* 5-day-old seedlings grown on +Fe (MS) plates were transferred to +Fe (MS) and –Fe (100µM FZ) plates for 72 hours. To study membrane damage, 300µM concentration of FerroZine was used (as described in figure legend). For expression analysis, seedlings were grown on +Fe (MS) solid media for 5 days. For the treatment, around 9-10 seedlings were transferred to 3ml of liquid +Fe (MS), and –Fe (300µM FZ), with or without 1µM Pep2 and treated for 24-72 hours depending upon need of the experiment (as described in figure legends). Seedlings were stained with PI solution (1 µg ml^−1^, dissolved in MilliQ water) for 2 min and rinsed in water. For propidium iodide (PI), 561-nm laser was used, and GFP/YFP was excited with a 488-nm laser, and emission spectra were collected at 600–650□nm, and 500–530□nm respectively. Imaging was performed in Z-stack mode with 1-μm step size on the Zeiss LSM710 confocal microscope. In each treatment, the Z-stack scan is processed to maximal Z-projection (Merge, PI and GFP/YFP). Scale bar: 100µm. Fiji® software Macro was used to quantify raw integrated density of the GFP/YFP signal.

### Perls’ and Perls’ DAB staining for histochemical detection of iron

5-day-old seedlings grown on +Fe (MS) were vacuum infiltrated with Perl’s staining solution (1% (v/v) HCl and 1% (w/v) K-ferrocyanide) for 25 minutes. The blue color intensity depicted the iron content. In order to stain both Fe^2+^ and Fe^3+^ forms, Perl’s stained seedlings were subjected to diaminobenzidine/DAB intensification. The seedlings were incubated for 1 hour in methanol containing 10□mm Na-azide and 0.3% (v/v) H2O2. The seedlings were washed twice with 100□mm Na phosphate buffer (pH 7.4) and incubated in DAB solution (0.025% (w/v) DAB and 0.005% (v/v) H2O2) for 2 minutes. The stained seedlings were washed three times with deionized water and imaged using NIKON ECLIPSE Ni U microscope.

### Rhizosphere acidification assay and pH quantification

7-day-old seedlings grown on +Fe (MS) plates were transferred to +Fe (MS) and –Fe (300µM FZ) plates for 72 hours. For plate assay, 9-10 seedlings were pooled together as one sample and placed onto rhizosphere acidification plates containing 0.7% agar and 0.006% (w/v) bromocresol purple (pH 6.5 adjusted with NaOH). After 72 hours the plates were scanned using Epson Perfection V600 at 1200□dpi resolution. For acidification capacity quantification assay, the seedlings were transferred to liquid medium supplemented with the pH indicator bromocresol purple (0.006%) for 2 days in 96-well plates. Proton extrusion capacity was analyzed by reading the absorption at 590 nm (A_590_) using an automated microplate reader.

### Histochemical GUS staining

To localize gene expression using GUS staining, 5-day-old seedlings grown on +Fe (MS) plates were transferred to +Fe (MS) and –Fe (300µM FZ) plates for 72 hours. The seedlings were vaccum-infiltrated in GUS staining solution containing 100□mM Na_2_HPO_4_, 2□mM□K_3_[Fe(CN)_6_], 2□mM□K_4_[Fe(CN)_6_], Triton X-100, 100□mM NaH_2_PO_4_, and 2□mM 5-bromo-4-chloro-3-indolyl glucuronide for 25 minutes. After GUS infiltration, the seedlings were incubated at 37°C for 1 hour, washed thrice with water and imaged using NIKON ECLIPSE Ni U microscope.

### ROS detection

For ROS detection with NBT (nitroblue tetrazolium) staining, the seedlings were kept in 20 mM Potassium phosphate buffer (pH 6.2) for 15 minutes and incubated in NBT staining solution (1 mM NBT solution in 20 mM Potassium phosphate buffer, pH 6.2) in dark for another 15 minutes. The stained seedlings were washed thrice with water. For clearing, seedlings were incubated in acidified methanol buffer (10 mL of methanol, 2 mL of HCl, 38 mL of water) at 57°C for 15 min and then in a basic solution (7% NaOH in 60% ethanol) for 15 min at room temperature. The seedlings were washed thrice and imaged using NIKON ECLIPSE Ni U microscope. The intensity of ROS accumulation was calculated using Fiji.

### Detection of Dead Cells with Evans Blue

To detect loss of cell viability using Evans blue staining, 5-day-old seedlings grown on +Fe (MS) plates were transferred to +Fe (MS) and –Fe (300µM FZ) plates for 72 hours. The seedlings were incubated in Evans solution (0.25% w/v Evans dye) for 20 minutes (Baker, C.J. and Mock, 1994). The stained seedlings were washed three times for 5 min each with distilled water and imaged using NIKON ECLIPSE Ni U microscope.

### RNA Extraction and qRT-PCR

5-day-old seedlings grown on +Fe solid medium were transferred to +Fe, –Fe, +Fe + Pep2 and –Fe + Pep2 in a 12-well plate and treated for 24 hours. Total RNA was extracted from around 60 roots using Plant RNeasy kit (Qiagen) following manufacturer’s protocol. 1µg of total RNA was reverse transcribed to cDNA using RevertAidTM First Strand cDNA Synthesis Kit (Thermofisher). qRT-PCR was performed with TB GreenTM Premix Ex using a Roche LightCycler 480 II. The gene expression level was normalized by TUBULIN as internal control. All the primers used for qRT-PCR analysis are listed in Table S1.

### Statistical analysis

Each experiment was repeated independently for at least 3 times with consistent results. Significance was calculated using Student’s t-test or one-way ANOVA, a post hoc Tukey HSD test. Different letters (a, b, c, and d) indicated significant differences. Error bars represented average ± SEM.

## ACKNOWLEDGEMENTS

We thank Jenny Russinova for providing *pepr1*, *pepr2*, *pepr1/2*, *pPEPR1:GUS:GFP/Col-0* and *pPEPR2:GUS:GFP/Col-0* seeds. We thank Janneke Balk for providing *bts*, Terry A. Long for *bts* and *pBTS:GUS*, Rumen Ivanov and Petra Bauer for *pIRT1:GUS*, Cyril Zipfel and Kalika Prasad for providing *rbohd* and *rbohf* seeds. We thank Vadthya Swapna and Swati Bhardwaj for their technical help in physiological experiments. We are grateful to all the members of SBS laboratory, Ajay Kumar Pandey, Amey Redkar and Vidha Srivastava for critical reading of the manuscript. SBS acknowledges intramural funding support from Indian Institute of Science Education and Research (IISER) Mohali. This work is supported by grant no. BT/PR51324/AGIII/103/1480/2023 from the Department of Biotechnology (DBT). SBS also acknowledges the Science and Engineering Research Board (SERB) for research funding (CRG/223 2022/003773). The SBS laboratory is also supported by the Indo□French Centre for the Promotion of Advanced Research (IFCPAR/ CEFIPRA) under project 68T06□1. DS acknowledges PhD fellowship from Council for Scientific and Industrial Research (CSIR), Govt. of India.

