## Supplementry Figures 1, 2, 3 and Table 1 for "An endogenous peptide PEP2 modulates Iron-deficiency signalling and root growth in Arabidopsis"

(a)

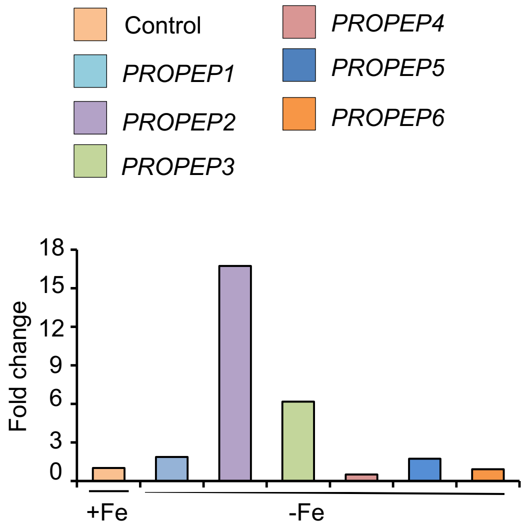


(b)

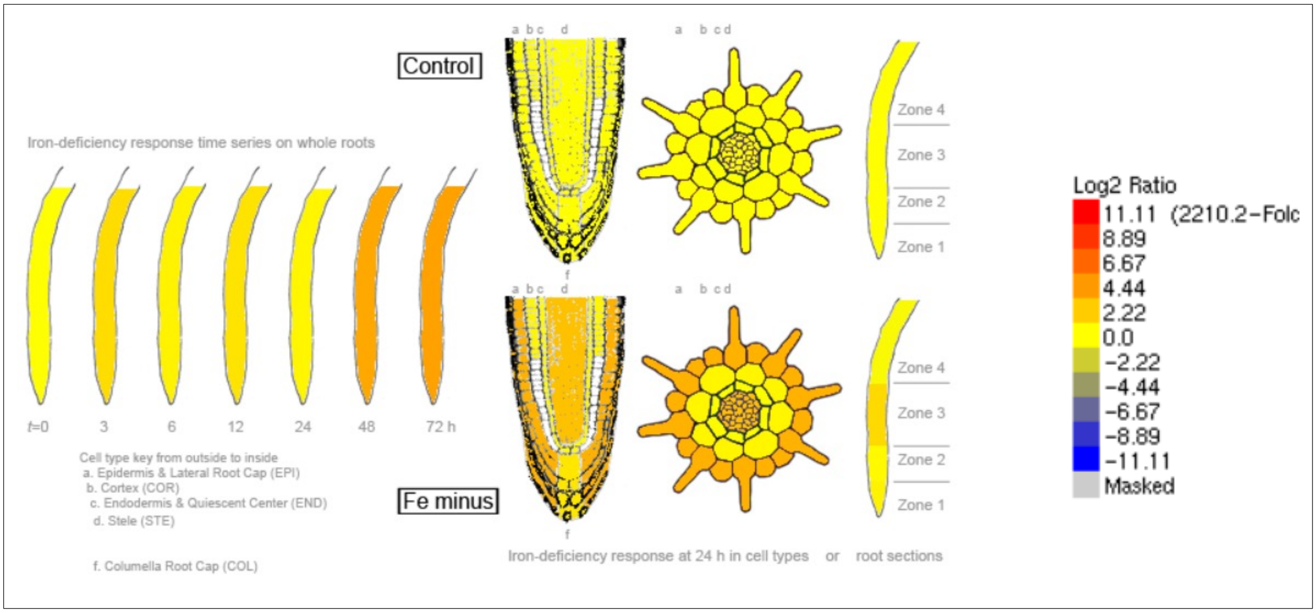


**Figure S1. *PROPEP2* is induced under Fe deficiency. (a-b)** Expression data from Arabidopsis eGFP browser. **(a)** Expression data of *PROPEP1-6* at 72 hours of iron deficiency with *PROPEP2* showing the highest expression. **(b)** Cell type or section type expression data generated by fluorescence-activated cell sorting of iron deprived roots.

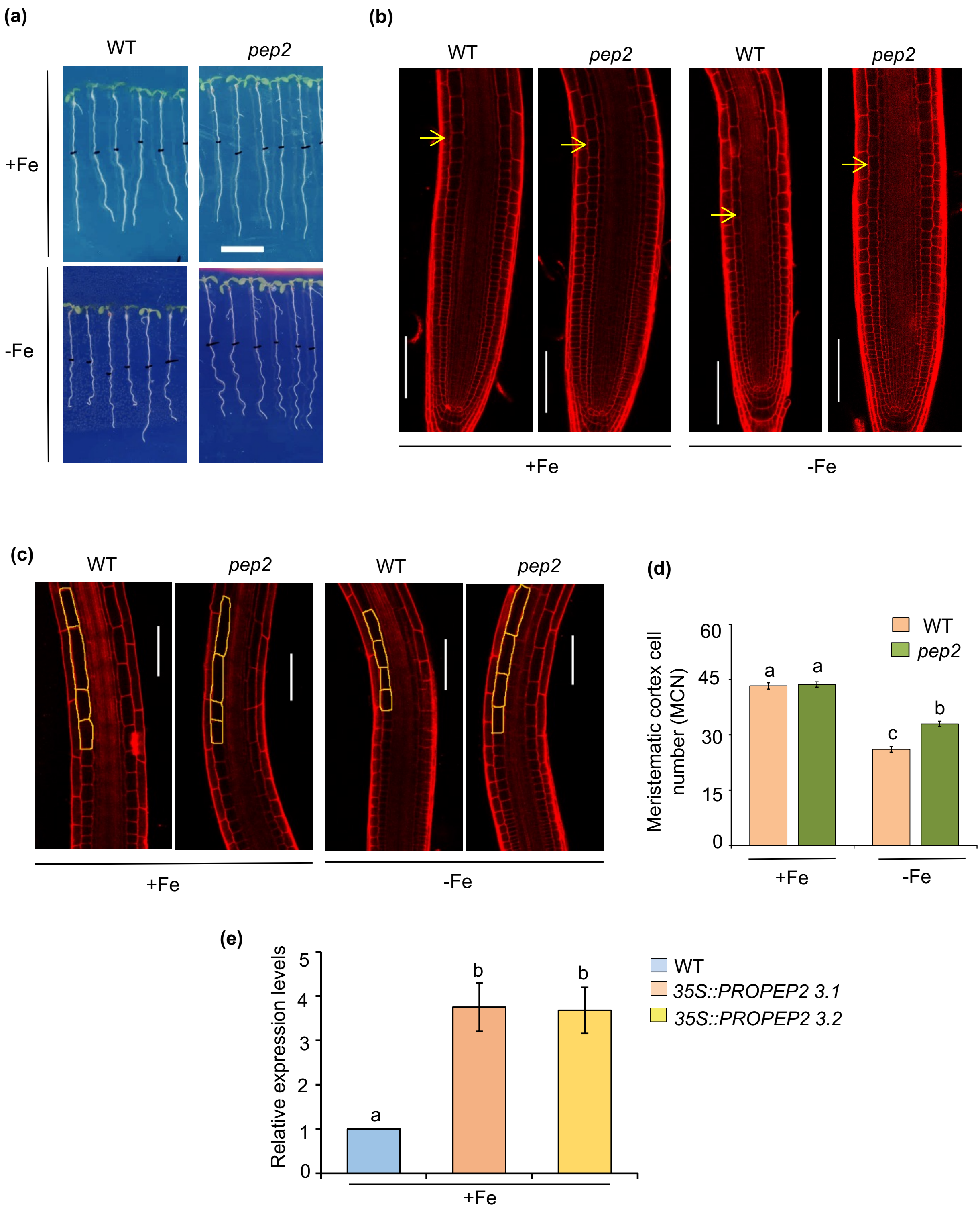


**Figure S2. *PEP2* negatively controls the meristem length and cell size in differentiation zone of root under Fe deficiency. (a-d)** Five days old seedlings of WT and *pep2* are transferred from +Fe media to +Fe and –Fe with 100μM Ferrozine for 72 hours and subjected to confocal microscopy. **(a)** Phenotypes of five days old seedlings that are transferred from +Fe media to +Fe and –Fe with 100μM Ferrozine for 72 hours. Scale bar: 1cm. **(b)** Confocal microscopy images of WT and *pep2* root tips. Yellow arrowheads indicate the cortex transition boundary (TB). Scale bar: 100μm. **(c)** Confocal microscopy images of elongation zone of WT and *pep2* root. Yellow line indicates cell boundary of elongating cell. Scale bar: 100μm. **(d)** Root meristem cortex cell number (MCN) of WT and *pep2* mutant. Error bars represent average ± standard errors (SE). Different letters (a, b, c, and d) indicate significant differences, as determined by one-way ANOVA and a post hoc Tukey Test (P≤0.05). **(e)** Expression levels of 35S::*PROPEP2*. Relative expression was determined by qRT-PCR in wild-type (WT) grown on +Fe media for 10 days. The gene expression level is normalized by TUB internal control. Data shown are an average of three biological replicates (n = 2 technical replicates). Each biological replicate consists of pooled RNA extracted from roots of ~90 seedlings. Error bars represent ± SEM. Significant difference by Student's t-test *(P ≤ 0.05).

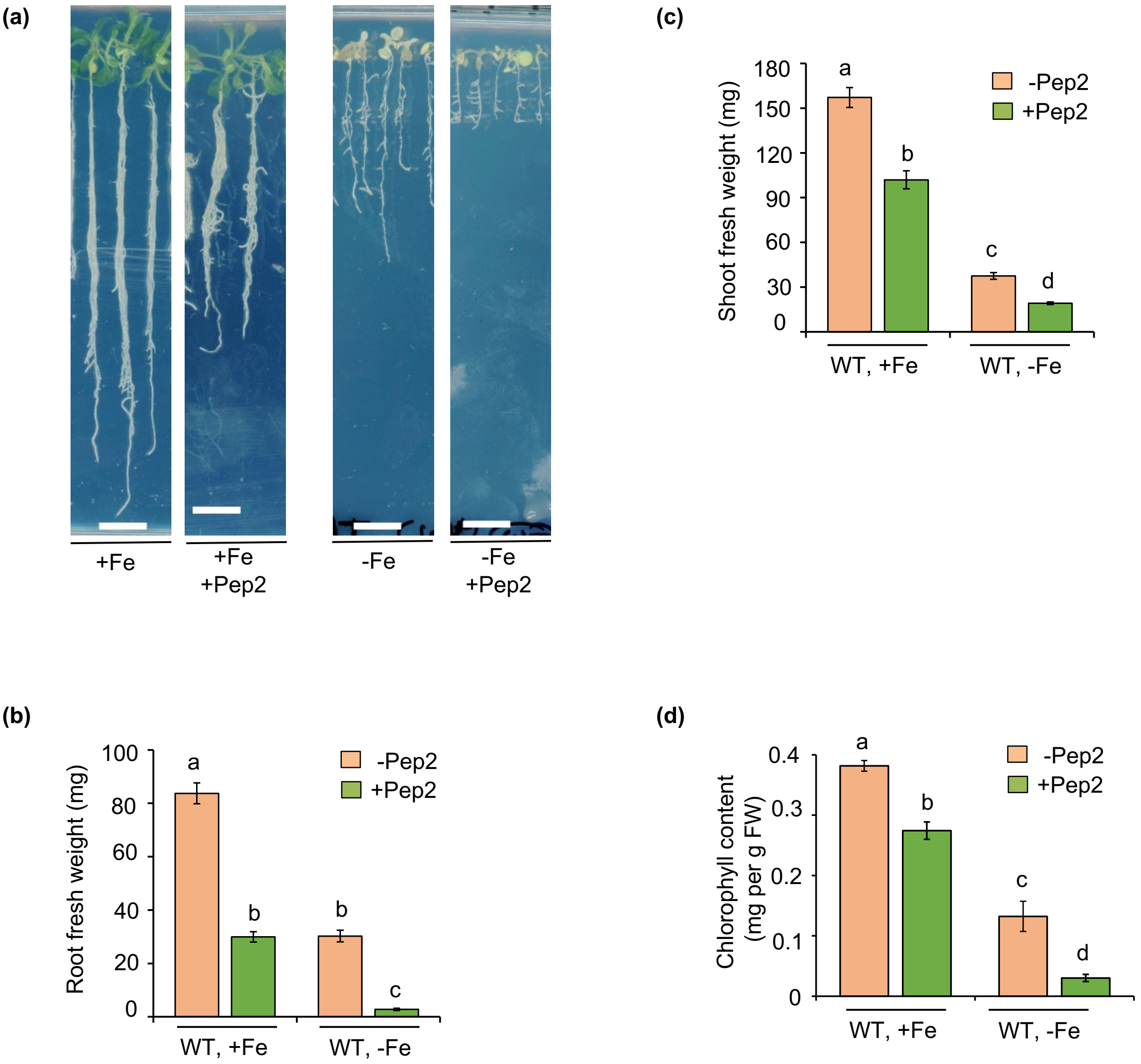


**Figure S3.** ***PEP2* negatively regulates plant growth and chlorophyll content under Fe deficiency**
**(a-d)** WT seedlings grown for five days on +Fe and then transferred for nine days to liquid +Fe and -Fe media with 100μM Ferrozine, with or without 50nM Pep2. **(b)** Root fresh weight (FW) of WT. **(c)** Shoot fresh weight (FW) of WT **(d)** The total chlorophyll concentration of the WT shoots. Error bars represent average ± standard errors (SE). Different letters (a, b, c,d and e) indicate significant differences, as determined by one-way ANOVA and a post hoc Tukey Test (P≤0.05). Scale bar: 1cm.

**Table S1.** Primers used in genotyping, qRT-PCR and cloning:

| | |
|---|---|
| *pep2* LP | GAGTCATCCAAAGCCACTAGC |
| *pep2* RP | TCTGGTCAATTTTGCTGTCTTG |
| *pepr1* LP | ACATCAGACGGACGTAAAACG |
| *pepr1* RP | TGCAATTAGGTGATCCGAAAC |
| *pepr2* LP | ACGGTGAACAAAATACGAACG |
| *pepr2* RP | TCTCAGATCTGCGGATAGCTC |
| LBb1.3 F | ATTTTGCCGATTTCGGAAC |
| qPEPR1 F | AACTAAAGGACTGTTATCAGTGGTAACG |
| qPEPR1 R | GGGCCTTCAATAAGCCATATTTT |
| qPEPR2 F | AAGAAGATGGCTTAATGCTG |
| qPEPR2 R | CAGTTGTGCCAGTAACAGTG |
| qPROPEP2 F | GTACGTAGTTTATTTTGGTTTCCTCATTT |
| qPROPEP2 R | GTCTTCAACAATTAGTTCCGATTCAA |
| qIRT1 F | GAATGTGGAAGCGAGTCAGCGA |
| qIRT1 R | GATCCCGGAGGCGAAACACTTA |
| qFRO2 F | GCCACATCTGCGTATCAAGTT |
| qFRO2 R | TCCCAAACAAGCTACGACCA |
| qRBOHD F | ACTCTCCGCTGATTCCAACG |
| qRBOHD R | ATCGCCGGAGACGTTATTCC |
| qRBOHF F | CTTGGCATTGGTGCAACTCC |
| qRBOHF R | TCTTTCGTCTTGGCGTGTCA |
| qBTS F | GCTCTGGCACAAGTCAATCA |
| qBTS R | CGTTCATCAAATGCCGATAA |
| qBTSL1 F | GGCAATGAAGATGGATTTGG |
| qBTSL1 R | TCATATGGAACCGTTGCTGA |
| qBTSL2 F | CGGGGCAGAATCCATCTTAT |
| qBTSL2 R | GTTGCAACAAGGAGCAAGAAG |
| pPROPEP2 FP | CACCCTATCTGTCTCTTTCAATTTCGTCG |
| pPROPEP2 FP+Sac1 | CACCGAGCTCCTATCTGTCTCTTTCAATTTCGTC |
| pPROPEP2 RP | TGAAATCCAATAGTTTGGTGAGTTATCG |
| pPROPEP2 RP+ Kpn1 | TTTGGTACCTGAAATCCAATAGTTTGGTGAGTTATCG |
| CDS PROPEP2 FP | CACCATGGAGAAATTAGATAAACGGAGG |
| CDS PROPEP2 RP | TTAATCCTCCTTATAAACTTGTATTGC |
